# Alveolar Epithelial Cell Loss of the Mitochondrial Regulator TFAM Drives Progressive Lung Fibrosis

**DOI:** 10.64898/2026.05.28.728065

**Authors:** Qianjiang Hu, Ugochi Onwuka, Nayra Cardenes, Madeline Packwood, Eileen L. Huang, Jian Shi, Maria Camila Melo-Narvaez, Puja Dutta, Zihe Zhou, Delphine Beaulieu, Byron Chuan, Pavan Suresh, Kevin Michael Redding, Laura-Marie Twardowski, Scarlett Varley, Ricardo H. Pineda, John Sembrat, Mara Lisa Grove Sullivan, Jonathan Franks, Simon C. Watkins, Claudette St. Croix, Corrine R. Kliment, Oliver Eickelberg, Mareike Lehmann, Marta Bueno, Brett A. Kaufman, Melanie Königshoff

## Abstract

Idiopathic pulmonary fibrosis (IPF) is characterized by failed alveolar epithelial repair and progressive fibrotic remodeling. Although aberrant reprogramming of alveolar type 2 (AT2) cells and accumulation of transitional AT2 states are increasing recognized as central features of IPF, the epithelial-intrinsic mechanisms that initiate these pathogenic states remain incompletely understood. Here, we identify mitochondrial transcription factor A (TFAM), a regulator of mitochondrial DNA maintenance, as a critical regulator of AT2 cell homeostasis. TFAM expression was reduced in AT2 cells from human IPF lungs. Inducible AT2 cell-specific *Tfam* deletion in mice caused spontaneous fibrotic remodeling and increased susceptibility to bleomycin-induced lung injury. TFAM-deficient AT2 cells acquired KRT8^+^ transitional and p21^+^ senescence-associated features before the onset of fibrotic transformation, accompanied by impaired oxidative phosphorylation, redox imbalance, mitochondrial superoxide accumulation, repression of mtDNA-encoded respiratory genes, and disrupted mitochondrial ultrastructure. TFAM-deficient AT2 cells developed a profibrotic secretory program that promoted extracellular matrix deposition and fibroblast activation. We further identified insulin-like growth factor-binding protein 2 (IGFBP2) as a secreted mediator induced in TFAM-deficient AT2 cells. IGFBP2 was elevated in AT2 cells in human IPF lung tissue and bronchoalveolar lavage fluid (BALF) from patients with IPF. IGFBP2 was detected in supernatants from fibrotic human precision-cut lung slices (hPCLS). IGFBP2 neutralization attenuated profibrotic remodeling in fibrotic hPCLS. Collectively, our findings identify TFAM-dependent mitochondrial homeostasis as an epithelial checkpoint linking AT2 cell-state stability to impaired epithelial-mesenchymal crosstalk driving pulmonary fibrosis.

## INTRODUCTION

Idiopathic Pulmonary Fibrosis (IPF) is a chronic, progressive interstitial lung disease with poor prognosis. It is characterized by relentless fibrotic scarring that ultimately leads to respiratory failure (1, 2). Despite a few available antifibrotic therapies, the disease remains irreversible and ultimately fatal for many patients (3). Increasing evidence indicates that defective epithelial repair is a central driver of disease pathogenesis (4, 5). In particular, lineage-tracing studies established that alveolar type 2 (AT2) cells function as alveolar progenitor cells in the adult lung, self-renewing and differentiating into alveolar type 1 (AT1) cells during homeostasis and after injury (6, 7), while their failure to maintain alveolar homeostasis is thought to initiate and sustain fibrotic remodeling (6). Defining the epithelial-intrinsic mechanisms that destabilize AT2 cell identity and compromise their reparative capacity is therefore critical to understanding IPF pathogenesis.

Recent single-cell studies in healthy and fibrotic lung tissues have identified aberrant epithelial transitional states as a hallmark of fibrotic lung remodeling (8–12). During healthy repair, AT2 cells can transiently adopt intermediate states prior to alveolar restoration. In the setting of pathological fibrosis, however, these programs become persistent and dysregulated. Keratin 8 (KRT8) positive transitional epithelial populations, together with senescence-associated epithelial programs, have emerged as key features of failed regeneration in both experimental fibrosis and human IPF (13–15). Mitochondrial dysfunction has been increasingly implicated in this process, particularly in the aging lung, where impaired mitochondrial homeostasis has been linked to epithelial vulnerability, epithelial senescence, defective regeneration, and fibrosis susceptibility (16–20). Nevertheless, the key mediator leading to mitochondrial failure within AT2 cells and whether this is sufficient to drive transitional and senescence-associated remodeling, independent of exogenous injury, remains unclear.

Transcription factor A, mitochondrial (TFAM), is a central regulator of mitochondrial DNA maintenance, transcription, and respiratory competence (21–25). Loss of TFAM disrupts mitochondrial homeostasis and can trigger downstream stress responses, including ROS accumulation, innate immune activation, inflammation, and metabolic reprogramming (26, 27). Although TFAM dysregulation has been implicated in fibrotic remodeling in other organs (28, 29), its role in the lung remains largely unexplored. In particular, whether AT2-specific TFAM deficiency is sufficient to trigger aberrant epithelial state transitions and spontaneous fibrotic remodeling, and how mitochondrial dysfunction in these cells reshape epithelial-mesenchymal communication, remain unknown.

To define the epithelial-intrinsic role of mitochondrial homeostasis in pulmonary fibrosis, we examined human IPF AT2 cells alongside an inducible AT2 specific *Tfam* deletion mouse model. Using these complementary approaches, we found that TFAM expression is reduced in AT2 cells from human IPF and that AT2 specific *Tfam* deletion is sufficient to induce progressive lung fibrosis. Before fibrosis develops, TFAM deficient AT2 cells acquire transitional and senescence-associated features alongside with mitochondrial dysfunction, indicating that TFAM is required to maintain epithelial state homeostasis. Mechanistically, TFAM deficient AT2 cells develop a broad profibrotic secretory program that promotes fibroblast activation and fibrotic remodeling. Within this altered epithelial secretome, insulin-like growth factor-binding protein 2 (IGFBP2) emerged as an induced downstream mediator, and analyses of human datasets and samples supported its clinical relevance and therapeutic potential. Together, these findings identify TFAM as an upstream epithelial regulator that links mitochondrial homeostasis to AT2 cell-fate stability and profibrotic epithelial-mesenchymal crosstalk in pulmonary fibrosis.

## RESULTS

### TFAM is reduced in AT2 cells in human IPF and experimental lung fibrosis

Mitochondrial dysfunction in AT2 cells is increasingly recognized as a feature of IPF and has been linked to defective epithelial repair and aberrant epithelial plasticity (16–18, 30–32). Consistent with this notion, analysis of the transcriptional data from Lung Genomics Research Consortium cohort (LGRC) (33) showed that *TFAM* transcript was significantly reduced in IPF lungs (n = 255) compared with donor controls (n = 137) and COPD lungs (n = 220) (Figure 1A). To test whether this reduction was associated with the alveolar epithelial compartment, we analyzed published scRNA-seq datasets from the IPF Cell Atlas (8–10), which revealed decreased *TFAM* expression in AT2 cells from IPF lungs relative to donor cells (Figure 1B). Similarly, pathway-level analysis of these cohorts demonstrated enrichment of mitochondrial dysfunction-related programs together with reduced oxidative phosphorylation activity in IPF AT2 cells, supporting a conserved shift toward impaired mitochondrial homeostasis in the diseased alveolar epithelium (Supplemental Figure 1). Immunofluorescence staining confirmed reduced TFAM protein abundance in LAMP3^+^ AT2 cells in IPF lungs compared with donor lungs (Figure 1C). Together, these findings demonstrate that TFAM expression is reduced in AT2 cells from human IPF patients.

**Figure 1.**
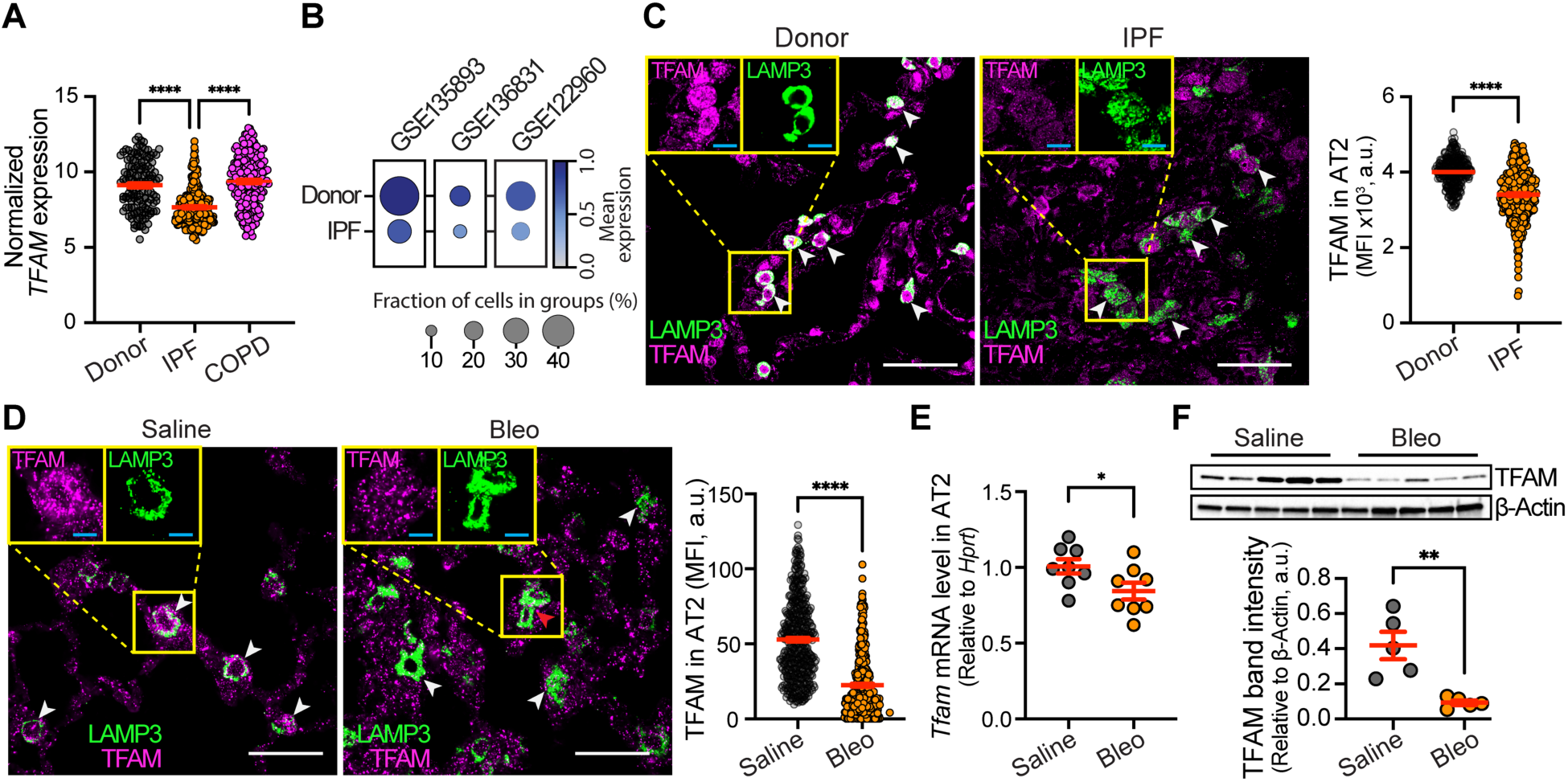
TFAM expression is decreased in AT2 cells in human IPF and experimental lung fibrosis. (A) Microarray analysis of the Lung Genomics Research Consortium cohort (LGRC; GSE47460) shows reduced *TFAM* transcript in IPF whole-lung tissue (n = 255) compared with donor controls (n = 137) and COPD lungs (n = 220). (B) IPF Cell Atlas scRNA-seq analysis shows decreased *TFAM* expression in AT2 cells from IPF lungs relative to donor lungs. (C) Representative immunofluorescence images and quantification show reduced TFAM protein in LAMP3^+^ AT2 cells in IPF relative to donor lungs (TFAM, magenta; LAMP3, green; arrowheads indicate AT2 cells). n = 6 per group; > 50 cells per individual were analyzed. Scale bars, 50 μm in overview images (white) and 10 μm in inset images (cyan). (D) Representative immunofluorescence images and quantification show reduced TFAM protein in LAMP3^+^ AT2 cells 14 days after bleomycin injury compared with saline controls. n = 6 per group; > 50 cells per mouse were analyzed. Scale bars, 50 μm in overview images (white) and 10 μm in inset images (cyan). (E) qPCR analysis of primary AT2 cells isolated 14 days after bleomycin injury shows reduced *Tfam* mRNA expression relative to saline-treated controls. n = 8 per group. (F) Immunoblot analysis of primary AT2 cells isolated 14 days after bleomycin injury confirms reduced TFAM protein abundance compared with saline-treated controls. n = 5 per group. Data are presented as mean ± SEM; a.u., arbitrary units; * *P* < 0.05, ** *P* < 0.01, **** *P* < 0.0001.

Consistent with the findings in human IPF, TFAM protein was also reduced in LAMP3^+^ AT2 cells *in situ* in the experimental lung fibrosis model of bleomycin-induced injury (Figure 1D). Primary AT2 cells isolated from bleomycin-treated mice showed significantly decreased *Tfam* mRNA and reduced TFAM protein abundance compared with saline-treated controls (Figure 1, E and F). Collectively, these human and experimental data identify reduced TFAM expression as a prominent feature of fibrotic AT2 cells and a potential driver of fibrogenesis.

### AT2-specific *Tfam* loss is sufficient to drive lung fibrosis

To explore whether AT2-intrinsic loss of TFAM is sufficient to drive fibrotic remodeling, we generated a tamoxifen-inducible, AT2 cell-specific conditional knockout model comprising wild-type controls (WT: *Sftpc*^+/+^; *Tfam*^fl/fl^), heterozygous mice (Het: *Sftpc*^CreERT2/+^; *Tfam*^fl/+^) and homozygous knockouts (KO: *Sftpc*^CreERT2/+^; *Tfam*^fl/fl^) (Figure 2A). Immunofluorescence analysis confirmed efficient, gene-dosage dependent reduction of TFAM mean fluorescence intensity (MFI) in LAMP3^+^ AT2 cells from Het and KO lungs compared with WT controls (Figure 2, B and C).

**Figure 2.**
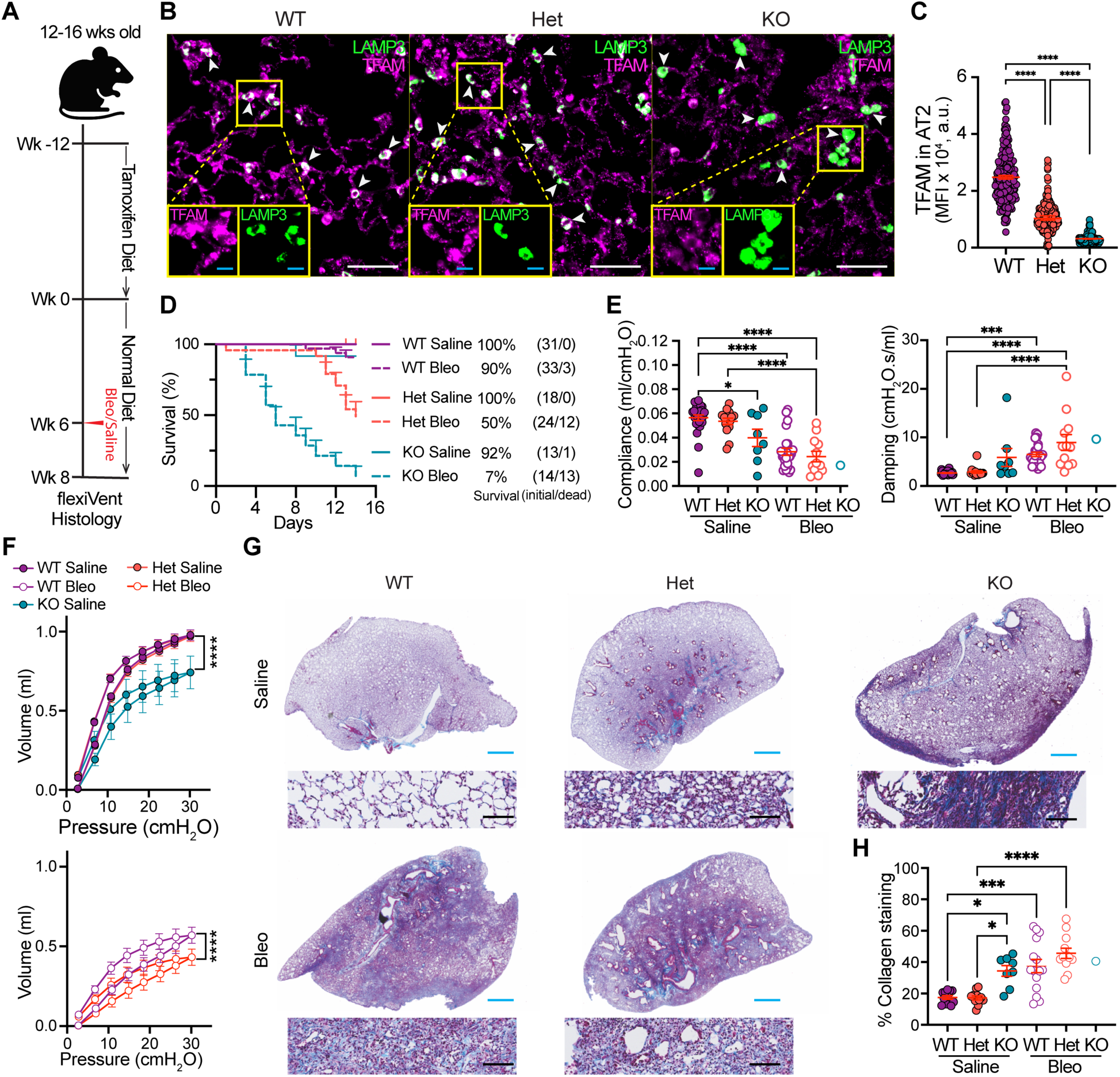
AT2-specific *Tfam* loss drives spontaneous and progressive lung fibrosis. (A) Schematic of the tamoxifen-inducible, AT2 cell-specific *Tfam* knockout model showing wild-type (WT: *Sftpc*^+/+^; *Tfam*^fl/fl^), heterozygous mice (Het: *Sftpc*^CreERT2/+^; *Tfam*^fl/+^) and homozygous knockouts (KO: *Sftpc*^CreERT2/+^; *Tfam*^fl/fl^) mice and timing of tamoxifen and bleomycin administration. (B, C) Representative immunofluorescence images and quantification show gene-dosage-dependent reduction of TFAM in LAMP3^+^ AT2 cells from WT, Het, and KO lungs. TFAM, magenta; LAMP3, green; arrowheads indicate AT2 cells. n = 6 per genotype; > 40 cells per mouse were analyzed. Scale bars, 50 μm in overview images (white) and 10 μm in insets (cyan). (D) Kaplan-Meier survival analysis shows reduced survival of KO mice and intermediate survival of Het mice following bleomycin injury. 2 U/kg bleomycin; n = 12-24 per group. (E) Lung function analysis shows reduced static compliance and increased tissue damping in saline-treated KO mice, with further impairment after bleomycin injury. Het mice show intermediate defects. n = 6-10 per group. (F) Pressure-volume loops confirm reduced lung compliance in Het and KO mice after saline or bleomycin treatment. (G, H) Masson’s trichrome staining and quantification show increased collagen deposition in saline-treated KO lungs and enhanced fibrosis after bleomycin injury. n ≥ 8 per group. Scale bars, 1 mm in whole lung images (cyan) and 100 μm in insets (black). Data are presented as mean ± SEM; * *P* < 0.05, *** *P* < 0.001, **** *P* < 0.0001.

We first examined whether loss of *Tfam* in AT2 cells alters susceptibility to fibrotic injury. We administered a single-dose bleomycin challenge (2 U/kg), a dose at which 10% of WT mice succumbed by day 14 (Figure 2D). Under these conditions, mice with AT2-specific TFAM deficiency showed increased susceptibility to bleomycin-induced injury. KO mice showed markedly reduced survival, with only 7% surviving to day 14, compared with 50% of the bleomycin-treated Het mice and 90% of bleomycin-treated WT mice (Figure 2D). Among surviving animals, bleomycin-treated Het mice showed more severe impairment in lung mechanics than bleomycin-treated WT mice, as reflected by lower static compliance, higher tissue damping, and altered pressure-volume loops (Figure 2, E and F). Histological analysis further demonstrated increased collagen deposition in Het lungs after bleomycin challenge, and quantitative image analysis confirmed increased collagen-positive area relative to WT bleomycin-treated controls (Figure 2, G and H). Altogether, these findings indicate that even a partial reduction of TFAM in AT2 cells increases vulnerability to injury-induced fibrotic remodeling.

Notably, even in the absence of exogenous injury, KO mice developed features of lung fibrosis as early as 8 weeks post-tamoxifen stimulation (Figure 2). Uninjured (saline-treated) KO mice showed reduced static compliance and increased tissue damping relative to WT controls, whereas respective Het mice did not show statistically significant changes at the same timepoint (Figure 2E). Consistent with these physiological abnormalities, pressure-volume loops demonstrated reduced lung compliance in the uninjured KO mice (Figure 2F). Histologically, Masson’s trichrome staining revealed spontaneous collagen accumulation in KO lungs at baseline, and quantification showed increased collagen-positive areas compared with WT controls (Figure 2, G and H). These findings were reproduced in an independent cohort and recapitulated using an alternative tamoxifen delivery strategy (Supplemental Figure 2), supporting the conclusion that AT2-specific TFAM deficiency is sufficient to initiate and drive fibrotic remodeling *in vivo*.

Because age is a major risk factor for IPF, and mitochondrial dysfunction is a hallmark of aging (1, 2, 34), we next evaluated the consequences of AT2-specific *Tfam* deletion in 12-month-old mice (Supplemental Figure 3A). Older mice harboring Tfam-deficient AT2 cells showed markedly reduced survival compared with their WT and Het counterparts with 10 out of 12 mice dead during the 10-week period after tamoxifen-diet completion (Supplemental Figure 3B). Surviving animals showed defects in lung mechanics, including reduced static compliance, increased tissue damping and abnormal pressure-volume loops (Supplemental Figure 3, C and D). Together, these data establish that AT2-specific *Tfam* loss not only enhances susceptibility to fibrotic injury but is sufficient to drive fibrosis development even in the absence of exogenous injury. Older mice exhibited an exaggerated phenotype.

### Tfam-deficiency drives transitional and senescence-associated AT2 cell states *in vivo*

To determine the mechanisms by which AT2-specific *Tfam* loss drives lung fibrosis, we examined lungs at an early time point, 1 week after completion of tamoxifen diet (Supplemental Figure 4A). At this timepoint, lung mechanics were preserved, and histological analysis showed no evidence of increased collagen deposition, supporting a pre-fibrotic window (Supplemental Figure 4B and C). Despite the absence of overt fibrosis, immunofluorescence analysis revealed an increased frequency of LAMP3^+^ AT2 cells coexpressing the transitional epithelial marker KRT8 in TFAM deficient lungs compared with WT controls, indicating early accumulation of a transitional AT2 population (Figure 3A). In parallel, *Tfam* KO lungs showed increased numbers of LAMP3^+^ AT2 cells expressing p21, consistent with induction of a senescence-associated epithelial phenotype (Figure 3B). Together, these findings indicate that AT2-intrinsic *Tfam* loss initiates epithelial reprogramming characterized by transitional and senescence-associated features, likely driving fibrotic remodeling.

**Figure 3.**
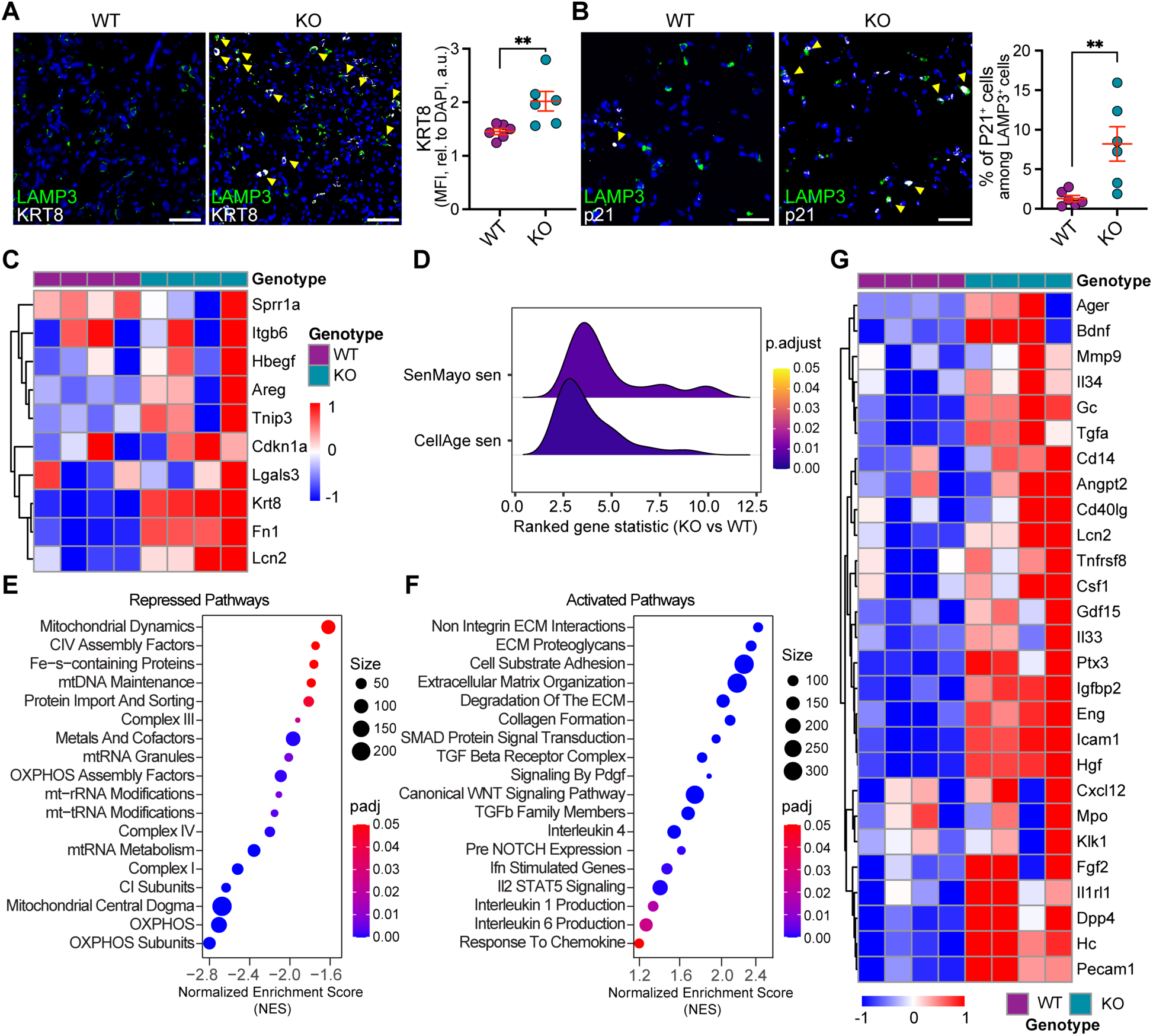
TFAM deficiency in AT2 cells promotest ransitional and senescence-like epithelial states at a pre-fibrotic stage. (A) Immunofluorescence staining and quantification show increased LAMP3^+^KRT8^+^ AT2 cells in AT2-*Tfam* KO lungs compared with WT controls (LAMP3, green; KRT8, gray; arrowheads indicate double-positive cells). n = 6 mice per group. Scale bars, 50 μm. (B) Immunofluorescence staining and quantification show increased LAMP3^+^p21^+^ AT2 cells in AT2-*Tfam* KO lungs (LAMP3, green; p21, gray; arrowheads indicate double-positive cells). n = 6 mice per group. Scale bars, 50 μm. (C) Bulk RNA-seq of whole lungs shows increased expression of KRT8^+^ transitional AT2 cell gene signatures in AT2-*Tfam* KO lungs. (D) Module-score analysis demonstrates enrichment of SenMayo and CellAge senescence-associated gene sets in AT2-*Tfam* KO lungs. (E, F) Gene set enrichment analysis shows repression of mitochondrial and oxidative phosphorylation pathways (E) and activation of extracellular matrix organization, TGF-β, and inflammatory signaling pathways (F) in AT2-*Tfam* KO lungs. (G) Heatmap showing upregulation of cytokine and chemokine genes, including senescence-associated secretory phenotype (SASP) components, in AT2-Tfam KO lungs. n = 4 mice per group for RNA-seq. Data are mean ± SEM where applicable; ** *P* < 0.01.

To define the transcriptional programs associated with these early epithelial changes, we performed bulk RNA-seq on whole lungs collected at the same pre-fibrotic time point (Supplemental Figure 5). Differential expression analysis confirmed reduced *Tfam* expression together with induction of epithelial stress and transitional markers, including *Krt8* (Supplemental Figure 5, A-C). TFAM deficient lungs exhibited increased expression of global *Krt8*^+^ and lipocalin 2 positive (*Lcn2*^+^) transitional AT2 cell signatures and enrichment of senescence-associated transcriptional programs, including SenMayo and CellAge gene sets (Figure 3, C and D) (35, 36). Gene set enrichment analysis further demonstrated repression of mitochondrial and oxidative phosphorylation pathways in TFAM-deficient lungs, together with activation of extracellular matrix related genes, TGF-β signaling, and inflammatory pathways (Figure 3, E and F). In addition, multiple cytokine and chemokine transcripts, including senescence-associated secretory phenotype (SASP)-related factors, were upregulated in TFAM-deficient lungs relative to WT controls (Figure 3G), and several established profibrotic genes were likewise increased (Supplemental Figure 5D). Together, these data show that loss of *Tfam* drives a pre-fibrotic epithelial program marked by mitochondrial suppression, transitional-state reprogramming, and senescence-associated signaling *in vivo*.

### TFAM loss drives cell-autonomous AT2 reprogramming and mitochondrial dysfunction

To determine whether these epithelial phenotypes reflect cell-intrinsic consequences of TFAM deficiency in AT2 cells, we established an *in vitro* deletion model by infecting primary mouse AT2 cells isolated from *Tfam*^fl/fl^ mice with control adenovirus (AdCre−) or adenoviral Cre recombinase (AdCre+) (Figure 4A). Efficient deletion was confirmed by significant suppression of *Tfam* transcript in AdCre+ cells (Figure 4B), together with a parallel reduction in mtDNA abundance relative to control cultures (Figure 4C).

**Figure 4.**
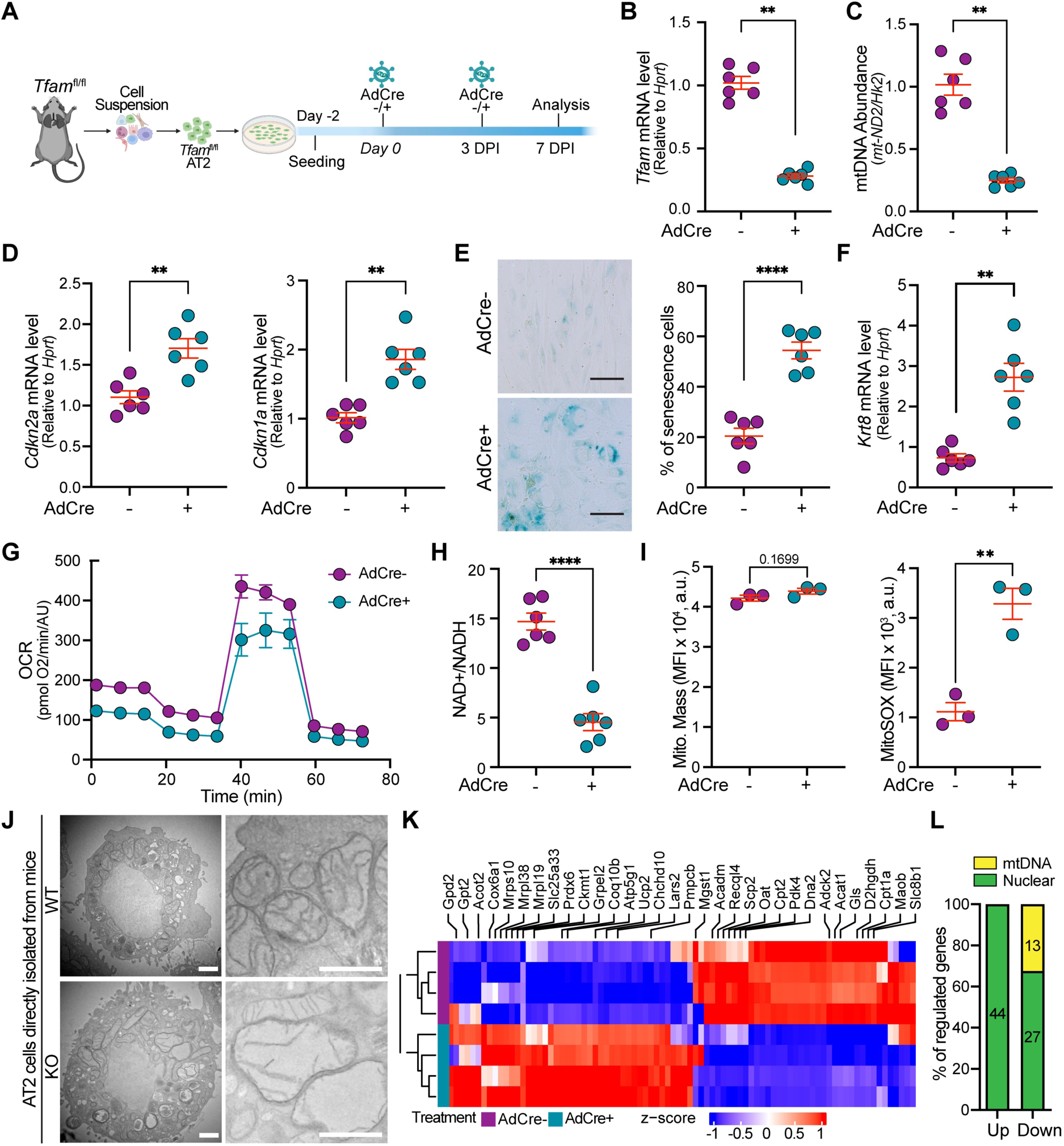
TFAM loss in primary AT2 cells induces mitochondrial dysfunction, cellular senescence, and a KRT8^+^ transitional epithelial state. (A) Experimental scheme. Primary AT2 cells from *Tfam*^fl/fl^ mice were infected with control adenovirus (AdCre−) or adenoviral Cre recombinase (AdCre+) and analyzed at 7 days post infection (DPI). (B, C) qPCR confirms reduced *Tfam* transcript and decreased mitochondrial DNA abundance, expressed as the mtDNA:nuclear DNA ratio, in AdCre+ cultures. (D, E) AdCre+ cultures show increased senescence-associated features, including elevated *Cdkn2a* and *Cdkn1a* expression and increased SA-β-gal^+^ cells. Scale bar, 100 μm. (F) qPCR shows increased *Krt8* expression in AdCre+ cultures. (G) Seahorse XF Cell Mito Stress Test shows reduced basal and maximal oxygen consumption rates (OCR) in TFAM*-*deficient AT2 cells relative to WT controls. (H) TFAM-deficient AT2 cells show a reduced NAD^+^/NADH ratio. (I) Flow cytometry analysis shows preserved mitochondrial mass measured by MitoTracker Deep Red but increased mitochondrial ROS measured by MitoSOX. (J) Transmission electron microscopy shows swollen mitochondria with collapsed cristae in AT2 cells isolated from *Tfam* KO mice, compared with intact cristae in AT2 cells isolated from WT mice. Scale bars, 1 μm. (K) Heatmap shows 84 differentially expressed MitoCarta 3.0 genes in AdCre+ versus AdCre− AT2 cells. (L) Genomic-origin analysis shows that all 44 upregulated MitoCarta-overlapping genes were nuclear encoded, whereas the 40 downregulated genes included 27 nuclear-encoded and 13 mtDNA-encoded protein-coding genes. For B-H, n = 6 per group; for I, n = 3 per group; for J, n = 4 mice per group; for K and L, n = 4 per group. Data are presented as mean ± SEM; ** *P* < 0.01, **** *P* < 0.0001.

Consistent with the pre-fibrotic epithelial phenotype observed *in vivo*, AdCre+ cultures increased senescence-associated features *in vitro*, as evidenced by increased expression of *Cdkn1a* and *Cdkn2a* transcript, increased SA-β-gal activity, and a higher proportion of p21^+^ cells (Figure 4, D and E; Supplemental Figure 6A). In parallel, TFAM-deficient cultures showed accumulation of KRT8^+^ transitional epithelial cells (Figure 4F; Supplemental Figure 6B). Together, these findings indicate that TFAM loss is sufficient to drive cell-autonomous senescence-associated and transitional-state remodeling in AT2 cells.

We next asked whether mitochondrial dysfunction was a defining feature of the TFAM-deficient AT2 cell state. Real-time live-cell oxygen consumption rate analysis by Seahorse XF analyzer showed that TFAM-deficient AT2 cells exhibited reduced basal and maximal oxygen consumption rates, consistent with impaired oxidative phosphorylation capacity (Figure 4G). In parallel, the NAD^+^/NADH ratio was significantly reduced, indicating disruption of mitochondrial redox homeostasis (Figure 4H). Flow cytometric analysis further showed that mitochondrial mass (MitoTracker) was preserved, whereas mitochondrial superoxide levels (MitoSox) were significantly increased following *Tfam* deletion (Figure 4I). TOM20 abundance remained comparable between control and *Tfam* deletion AT2 cells (Supplemental Figure 6C). These data indicate that TFAM deficiency impairs mitochondrial function, increases reactive oxygen species production without changes in mitochondria mass.

Notably, these functional defects were mirrored by profound changes in mitochondrial ultrastructure. In contrast to the intact architecture observed in WT AT2 cells, TFAM-deficient AT2 cells displayed marked mitochondrial abnormalities, characterized by significant swelling and the presence of disrupted or collapsed cristae (Figure 4J).

To determine how TFAM deficiency reshapes mitochondrial gene programs in AT2 cells, we intersected bulk RNA-seq differentially expressed genes (DEGs) from AdCre+ versus AdCre− cultures with the MitoCarta 3.0 gene set (37). This analysis identified 84 mitochondria-associated DEGs (Figure 4K). Stratification of these genes by genomic origin revealed that all 44 upregulated genes were nuclear-encoded, whereas the 40 downregulated genes included 27 nuclear-encoded genes together with all 13 mtDNA-encoded protein-coding genes (Figure 4L), suggesting that TFAM loss disrupts not only mtDNA-derived respiratory gene expression but also a broad nuclear-encoded mitochondrial gene network. Collectively, these data show that TFAM loss causes profound mitochondrial dysfunction in AT2 cells and is coupled to senescence-associated and transitional epithelial reprogramming.

### TFAM-deficient AT2 cells acquire a profibrotic secretory program that remodels the lung microenvironment

Given the strong cell-intrinsic AT2 phenotype induced by TFAM loss, we next asked how TFAM-deficient AT2 cells might influence the surrounding microenvironment and contribute to structural lung remodeling. To address this, we applied weighted gene co-expression network analysis (WGCNA) to bulk RNA-seq data from control and *Tfam*-deleted primary AT2 cells to identify coordinated gene modules altered after TFAM loss (Supplemental Figure 7). This analysis identified 2 contrasting modules strongly associated with genotype. A turquoise module was negatively correlated with *Tfam* knockout status and enriched for mitochondrial related genes, consistent with suppression of an expected mitochondrial homeostatic program in TFAM-deficient AT2 cells (Supplemental Figure 7 B and C). In contrast, a blue module was strongly positively correlated with *Tfam* knockout status, and its eigengene was significantly increased in *Tfam*-deleted AT2 cells, indicating that this gene module is activated after *Tfam* loss. (Figure 5A; Supplemental Figure 7). Functional enrichment analysis of this module revealed pathways related to cytokine production, inflammatory signaling, cell communication, extracellular matrix organization, and lung fibrosis, consistent with acquisition of a profibrotic secretory program (Figure 5B). Within blue module, genes with high module membership and strong association with *Tfam* knockout included transcripts associated with *Krt8*^+^ alveolar differentiation intermediate (ADI) cells (14) together with multiple profibrotic and inflammatory mediators, supporting the emergence of a stress-associated epithelial state with profibrotic signaling potential after *Tfam* loss (Figure 5C). Consistent with these network-level changes, gene-level analysis further showed increased expression of cytokine/chemokine, inflammatory mediator, and profibrotic transcripts in TFAM-deficient AT2 cells (Supplemental Figure 8). Together, these data indicate that TFAM-deficient AT2 cells are transcriptionally primed to remodel the surrounding microenvironment through mitochondrial dysfunction-associated secretory reprogramming.

**Figure 5.**
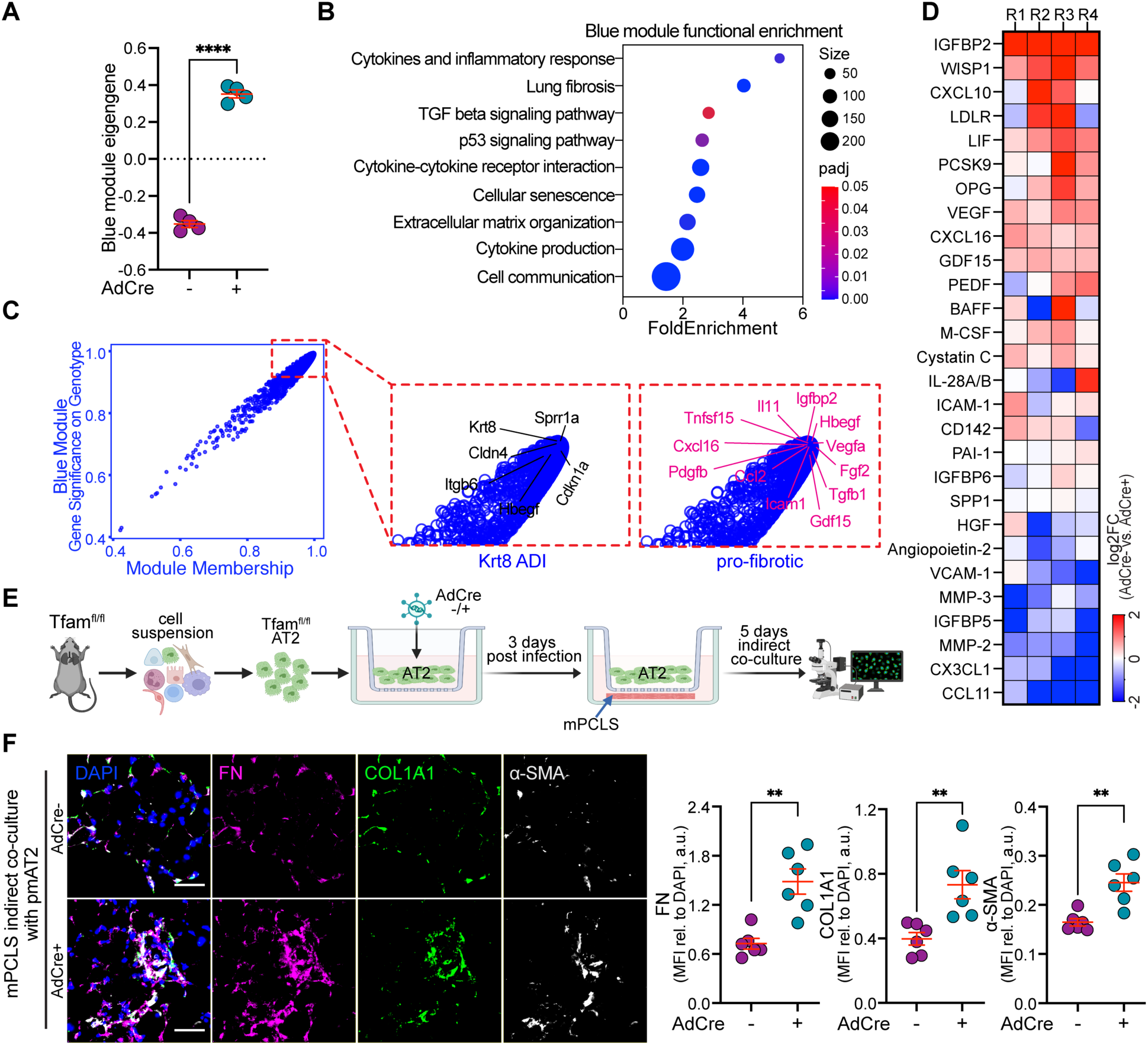
TFAM-deficient AT2 cells activate a profibrotic secretory program and promote paracrine tissue remodeling. Bulk RNA-seq data from control (AdCre−) and *Tfam*-deleted (AdCre+) primary AT2 cells were analyzed by weighted gene co-expression network analysis (WGCNA). (A) Blue module eigengene plot showing increased activity of the KO-associated blue WGCNA module in AdCre+ AT2 cells relative to AdCre− controls. Each dot represents one biological replicate. (B) Functional enrichment analysis of blue module genes showing enrichment for pathways related to cell communication, cytokine production and inflammatory responses, extracellular matrix organization, cellular senescence, TGF-β signaling, and lung fibrosis. (C) Scatter plot showing the relationship between module membership and gene significance for genotype within the blue module. Representative genes associated with Krt8^+^ alveolar differentiation intermediate (ADI) and profibrotic programs are highlighted. (D) Cytokine array of AT2-derived conditioned media collected from control (AdCre−) and *Tfam*-deleted (AdCre+) AT2 cells, showed as heatmap of log2 fold change (AdCre+ versus AdCre−). n = 4 per group. (E) Schematic of the indirect co-culture experiment. Created in BioRender. Primary mouse AT2 cells isolated from *Tfam*^fl/fl^ mice were infected with control adenovirus (AdCre−) or adenoviral Cre recombinase (AdCre+) to induce *Tfam* deletion and then indirectly co-cultured with mPCLS. (F) Representative immunofluorescence images of mPCLS indirectly co-cultured with AdCre− or AdCre+ AT2 cells and stained for fibronectin (FN, magenta), COL1A1 (green), and α-SMA (gray), with DAPI (blue), show increased extracellular matrix deposition and myofibroblast marker expression in slices exposed to TFAM-deficient AT2 cells. Quantification of FN, COL1A1, and α-SMA mean fluorescence intensity (MFI; arbitrary units) is shown at right. n = 6 per group. Scale bar, 50 μm. Data are presented as mean ± SEM; ** *P* < 0.01, **** *P* < 0.0001.

We next performed cytokine analysis of conditioned media from TFAM-deficient and control AT2 cells (Figure 5D). Notably, multiple profibrotic and senescence-associated mediators were upregulated, including factors previously implicated in profibrotic signaling, such as WISP1 (38, 39), and senescence-associated secretory phenotype (SASP) related factors GDF15 and IGFBP2 (40). To test whether these secreted factors were functionally active, we used an indirect co-culture system in which TFAM-deficient (or control) AT2 cells were cultured with mouse precision-cut lung slices (mPCLS) (Figure 5E). Indirect co-culture with TFAM-deficient AT2 cells increased fibronectin, collagen I A1 (COL1A1), and α-smooth muscle actin (SMA) protein expression in mPCLS compared with co-culture with control AT2 cells, indicating enhanced extracellular matrix deposition and myofibroblast-associated remodeling (Figure 5F). In support, conditioned medium from TFAM-deficient AT2 cells accelerated lung fibroblast migration in wound-scratch assays, demonstrating a paracrine effect on mesenchymal cell behavior (Supplemental Figure 9). Together, these data show that TFAM-deficient AT2 cells acquire a biologically active profibrotic secretory program that promotes remodeling of the lung microenvironment.

### IGFBP2 contributes to fibroblast activation downstream of TFAM-deficient AT2 cells and is elevated in human IPF

IGFBP2, a SASP factor previously reported to be elevated in serum and sputum from patients with IPF (40–42), was among the most strongly increased proteins in conditioned media from TFAM-deficient AT2 cells (Figure 5D). Consistent with this finding, *Igfbp2* expression was increased at the transcript level in both *Tfam*-deleted AT2 cells and whole lungs from AT2-specific *Tfam* knockout mice (Supplemental Figure 10, A and B). Immunofluorescence staining further confirmed increased IGFBP2 expression in AT2 cells in *Tfam* KO lungs compared with WT controls (Figure 6A). In AT2 lineage-tracing *Tfam* knockout mice, GFP-labeled AT2-lineage cells co-expressed KRT8 and IGFBP2, indicating that disease-associated transitional AT2-lineage cells are one source of IGFBP2 in the fibrotic lung (Supplemental Figure 10C). These findings suggest IGFBP2 as an induced component of the broad TFAM-deficient epithelial secretory program and nominate it as a candidate mediator of downstream profibrotic signaling.

**Figure 6.**
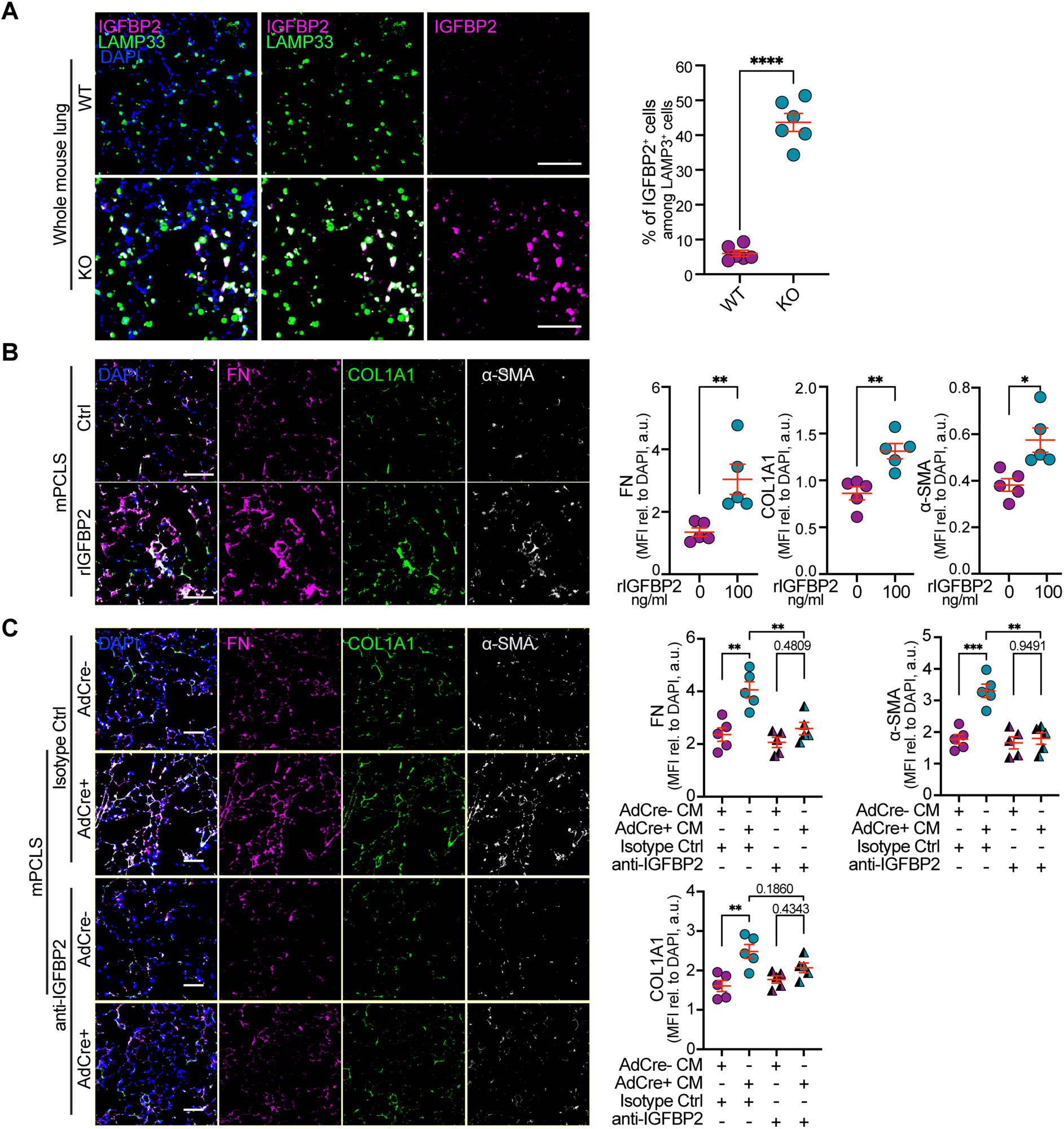
TFAM-deficient derived IGFBP2 promotes fibrotic remodeling in *ex vivo* lung tissue. (A) Immunofluorescence staining and quantification show increased IGFBP2 expression in LAMP3^+^ AT2 cells in AT2-*Tfam* KO mouse lungs compared with WT controls. n = 6 per group. Scale bar, 100 μm. (B) mPCLS treated with recombinant IGFBP2 (rIGFBP2) showe enhanced fibrotic remodeling, as assessed by immunofluorescence staining for fibronectin (FN, magenta), COL1A1 (orange), α-SMA (white), and DAPI (blue). n = 5 per group. Scale bar, 100 μm. (C) Representative immunofluorescence images of mPCLS treated with conditioned medium from Tfam-deleted AdCre+ AT2 cells supplemented with an IGFBP2-neutralizing antibody (anti-IGFBP2) or isotype control. Staining for FN, COL1A1, α-SMA, and DAPI shows that IGFBP2 blockade attenuates extracellular matrix deposition and myofibroblast marker expression induced by conditioned medium from TFAM-deficient AT2 cells. n = 5 per group. Scale bar, 100 μm. Data are presented as mean ± SEM; * *P* < 0.05, ** *P* < 0.01, *** *P* < 0.001, **** *P* < 0.0001.

We thus investigated whether IGFBP2 functionally contributes to the profibrotic activity of TFAM-deficient AT2 cells. Indeed, treatment of mPCLS with recombinant IGFBP2 increased fibronectin, COL1A1, and α-SMA (Figure 6B). Conversely, in mPCLS exposed to conditioned medium from TFAM-deficient AT2 cells, IGFBP2 neutralizing antibody (anti-IGFBP2) reduced fibronectin and α-SMA expression and partially reduced COL1A1 expression relative to isotype-treated controls (Figure 6C). Together, these data indicate that IGFBP2 is a functional component of the TFAM-deficient AT2 secretome, contributing to fibroblast activation and *ex vivo* tissue remodeling.

Extending these observations to human disease, analysis of the transcriptional data from LGRC cohort (33) demonstrated significantly increased IGFBP2 transcript abundance in whole-lung tissue from patients with IPF compared with donor controls and COPD lungs (Figure 7A). Analysis of multiple published single-cell RNA-seq datasets further showed increased IGFBP2 expression in AT2 cells from IPF lungs relative to donor controls (Figure 7B). Moreover, increased IGFBP2 protein was detected in bronchoalveolar lavage fluid (BALF) from IPF patients compared with those from non-ILD controls (Figure 7C). Consistently, immunofluorescence staining of human lung tissue showed increased IGFBP2 expression predominantly in HTII-280^+^ AT2 cells in IPF compared with donor lungs (Figure 7D). Together, these data demonstrate that IGFBP2 is elevated in human IPF lungs, enriched in IPF AT2 cells, and increased in BALF, supporting its relevance as a disease-associated epithelial secretory mediator.

**Figure 7.**
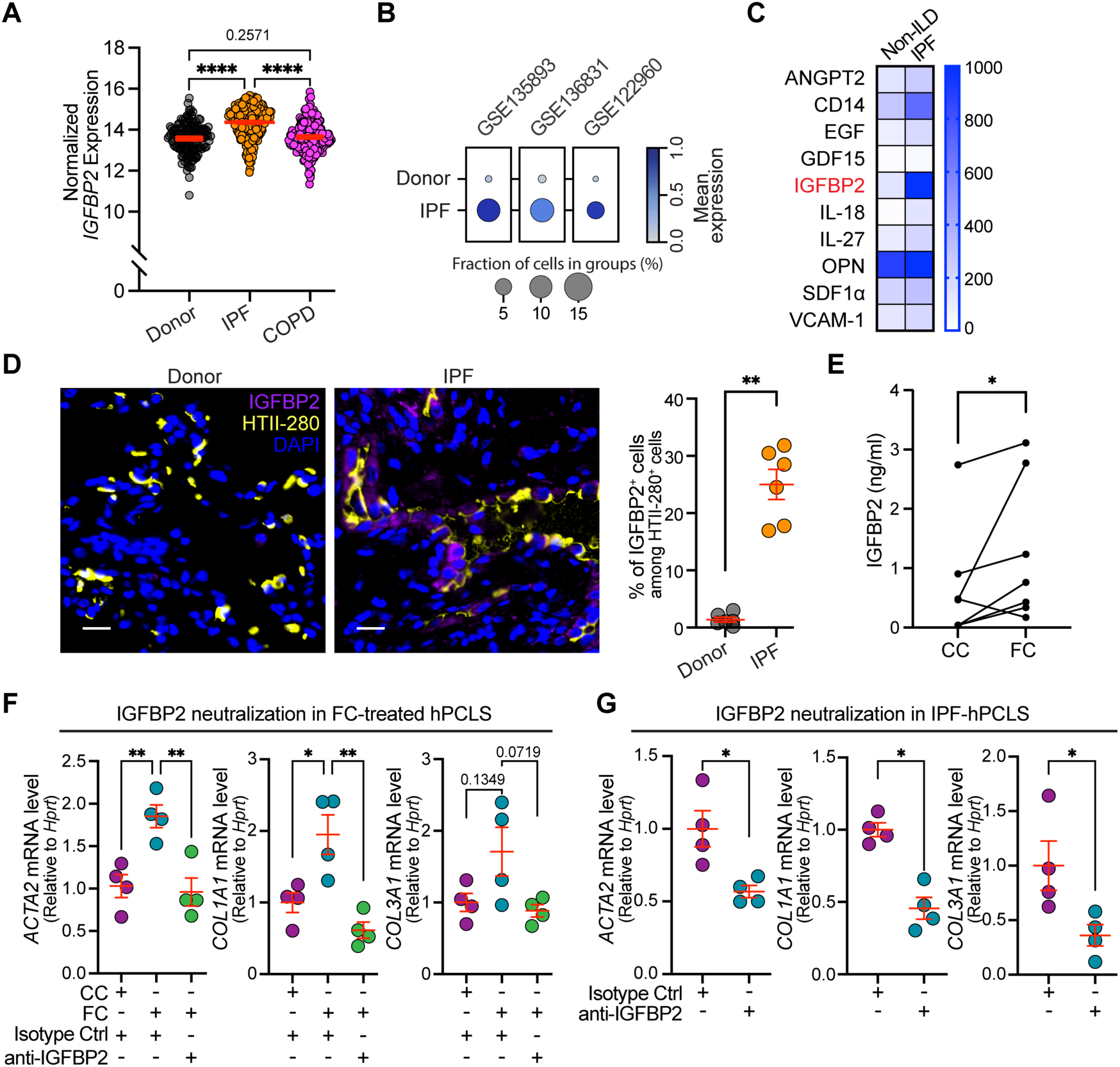
IGFBP2 is a secreted and targetable profibrotic mediator in human pulmonary fibrosis. (A) Microarray analysis of the Lung Genomics Research Consortium cohort (LGRC; GSE47460) shows increased IGFBP2 transcript in IPF whole-lung tissue (n = 255) compared with donor controls (n = 137) and COPD lungs (n = 220). (B) Single-cell RNA-seq analysis of human lung datasets (GSE135893, GSE136831, and GSE122960) show increased IGFBP2 expression in AT2 cells from IPF lungs relative to donor controls. (C) Cytokine array analysis of bronchoalveolar lavage fluid (BALF) identifies increased cytokines, including IGFBP2, in BALF from patients with IPF compared with non-ILD controls; 5 BALF samples were pooled per group. (D) Representative immunofluorescence staining of human lung sections showing increased IGFBP2 expression in HTII-280^+^ AT2 cells in IPF compared with donor lungs. HTII-280, yellow; IGFBP2, magenta; DAPI, blue. n=6 per group. Scale bar, 20 μm. (E) ELISA of conditioned medium from human precision-cut lung slices (hPCLS) treated with control cocktail (CC) or fibrotic cocktail (FC) shows increased IGFBP2 secretion following FC treatment. Each connected pair represents hPCLS derived from the same donor lung; n = 7. (F) qPCR analysis of hPCLS treated with CC or FC, with or without an IGFBP2-neutralizing antibody (anti-IGFBP2) or isotype control, shows that IGFBP2 neutralization attenuates FC-induced *ACTA2*, *COL1A1*, and *COL3A1* expression; n = 4. (G) qPCR analysis of 4 independent paired IPF hPCLS samples generated from one explanted lung and treated with an IGFBP2-neutralizing antibody or isotype control shows reduced *ACTA2*, *COL1A1*, and *COL3A1* expression following IGFBP2 blockade. Data are presented as mean ± SEM; * *P* < 0.05, ** *P* < 0.01, *** *P* < 0.001, **** *P* < 0.0001.

To test whether IGFBP2 contributes to profibrotic remodeling in human lung tissue, we next used an *ex vivo* human precision-cut lung slices (hPCLS) modes of pulmonary fibrosis. We found that fibrotic cocktail (FC)-treated hPCLS secreted higher IGFBP2 levels than control cocktail (CC)-treated hPCLS (Figure 7E). Importantly, IGFBP2 neutralizing antibody (anti-IGFBP2) attenuated FC-induced expression of *ACTA2*, *COL1A1*, and *COL3A1* (Figure 7F). We further examined hPCLS from one explanted IPF lung and found that treatment with the anti-IGFBP2 reduced *ACTA2*, *COL1A1*, and *COL3A1* expression relative to isotype-treated controls (Figure 7G). Overall, these data support IGFBP2 as an AT2 cell-derived profibrotic mediator linked to epithelial mitochondrial dysfunction and identify IGFBP2 neutralization as a potential therapeutic strategy for human pulmonary fibrosis.

## DISCUSSION

In this study, we identify TFAM-dependent mitochondrial maintenance as a critical checkpoint that preserves AT2 cell homeostasis and prevents fibrotic lung remodeling. We show that expression of TFAM is reduced in AT2 cells in human IPF lung tissue as well as mouse experimental lung fibrosis. AT2-specific *Tfam* deletion in mice is sufficient to cause a decline in lung function and progressive fibrotic remodeling *in vivo*, even in the absence of exogenous injury. TFAM-deficient AT2 cells acquire senescence-associated and transitional-state features, along with profound mitochondrial dysfunction, before the onset of tissue fibrosis, indicating that loss of mitochondrial integrity precedes and potentially initiates aberrant epithelial remodeling. TFAM-deficient AT2 cells develop a profibrotic secretory program that induces fibroblast activation and lung remodeling. Importantly, we identified IGFBP2 as an emerging downstream mediator of mitochondrial reprogramming. These findings thus position TFAM as an upstream regulator linking epithelial mitochondrial homeostasis, AT2 cell-state stability, and profibrotic epithelial-mesenchymal crosstalk in pulmonary fibrosis. IGFBP2 may serve as a therapeutically valid target to attenuate lung fibrosis.

Over the last decade, AT2 cell reprogramming has emerged as a critical feature of pulmonary fibrosis (43–45). More recently, human single-cell studies have reshaped our understanding of IPF by showing that fibrotic lungs undergo profound epithelial remodeling, including loss of normal alveolar epithelial identity and emergence of aberrant transitional or basaloid epithelial populations (8–10, 13). These single cell atlases identified rare senescent epithelial cells (8), KRT5-/KRT17+ aberrant basaloid cells localized near fibroblast foci (9, 10), and transitional alveolar epithelial cells with ineffective AT1 differentiation (13). Similarly, mouse injury-repair studies identified Krt8+ alveolar intermediate states that emerge during AT2-to-AT1 regeneration but persist in fibrotic remodeling, with related KRT8-associated transitional programs also detected in human fibrosis (13, 14, 46). Together, these studies provided unprecedented detail into AT2 cell reprogramming and further corroborated aberrant alveolar epithelial cell states as a central feature of pulmonary fibrosis. However, the epithelial-intrinsic mechanisms that initiate or sustain these pathological states remain largely unresolved. Here, our findings add TFAM downregulation and impaired mitochondrial homeostatic programs to this disease-associated epithelial landscape, and raise the possibility that mitochondrial dysfunction represents an upstream mechanism contributing to AT2 state failure in IPF.

A key advance of our study is the demonstration that an AT2-intrinsic mitochondrial defect is sufficient to initiate and drive fibrotic remodeling. Prior work has shown that AT2 dysfunction can promote fibrosis through multiple mechanisms, including epithelial senescence, impaired alveolar regeneration with mechanical tension-induced TGF-β signaling, and persistence of aberrant intermediate epithelial states that reinforce fibroblast activation and inflammatory niche remodeling (13, 47–50). However, most of these epithelial states have been characterized in the setting of exogenous injury or established fibrotic disease (13, 14, 46, 51). Thus, it remains unclear whether they represent primary epithelial defects capable of initiating fibrosis or secondary consequences of a profibrotic multicellular cascade. Our novel inducible genetic model addresses this gap by demonstrating that AT2-specific deletion of a major mitochondrial regulator (*Tfam*), without bleomycin or other exogenous injury, is sufficient to cause progressive lung fibrosis. Moreover, TFAM-deficient AT2 cells acquired transitional and senescence-associated features before detectable fibrosis developed, supporting the concept that aberrant epithelial-state remodeling can precede, and initiate mesenchymal remodeling. These findings position TFAM-dependent mitochondrial integrity as an epithelial-intrinsic determinant of fibrotic susceptibility and suggest that mitochondrial genome failure can serve as an initiating defect in the adult alveolar epithelium. Future studies will be required to determine the relative contributions of direct AT2-derived profibrotic signaling and its secondary amplification through immune cell recruitment, cell-cell, and cell-ECM interaction.

The spontaneous fibrosis observed after AT2-specific *Tfam* deletion should be considered alongside other epithelial-intrinsic models of progressive fibrotic lung disease. Surfactant protein mutations are clinically relevant drivers of IPF and mutant *Sftpc* models have shown that defects in AT2 proteostasis, impairment of macroautophagy, alongside ER stress can drive spontaneous inflammation followed by fibrotic remodeling, supporting AT2 quality-control failure as a proximal driver of lung fibrosis (52–54). Notably, the mouse model of mutant *Sftpc*^I73T^ expression has recently been shown to exhibit impaired AMPK signaling and AT2 metabolism, leading to accumulation of KRT8^+^ transitional epithelial cells (54). AMPK regulates multiple metabolic enzymes and mitochondrial quality-control pathways and acts upstream of TFAM (55, 56). Interestingly, metformin has been shown to activate AMPK signaling and increased TFAM expression in fibrotic fibroblasts leading to a reduction in ECM markers (57). Impaired AMPK signaling together with TFAM deficiency may further compromise AT2 cell resilience and contribute to the emergence of dysfunctional transitional AT2 cell states in IPF. TFAM-deficient AT2 cells further show upregulation of senescence marker and targeting cellular senescence has been demonstrated to attenuate fibrosis (34, 49, 50, 58). In line with this concept, AT2-specific *Sin3a* deletion has showed that p53-dependent epithelial senescence alone can drive spontaneous progressive pulmonary fibrosis (47). In another AT2-intrinsic model, loss of CDC42, a Rho GTPase required for actin filament polymerization, in AT2 cells impaired AT2-to-AT1 differentiation and promoted progressive fibrosis through sustained mechanical tension and AT2 TGF-β activation. Notably, the fibrotic phenotype was accelerated by pneumonectomy, whereas spontaneous fibrosis in untreated *Cdc42*-null mice was observed only in aged animals (48). Our findings further support a critical cell-intrinsic role for AT2 cells in fibrotic remodeling. Using an adult AT2-specific model of TFAM-dependent mitochondrial disruption, we identify mitochondrial dysfunction, a prominent hallmark of aging, as an AT2 cell-intrinsic driver of failed epithelial resilience and aberrant repair, rather than merely a secondary consequence of lung injury (59).

We found a prominent increase of transitional AT2 cells upon loss of TFAM expression. Prior studies identified KRT8^+^ ADI, pre-alveolar type-1 transitional cell state (PATS), and damage-associated transient progenitors (DATPs) as AT2-derived states that emerge during injury repair and are enriched for stress-response programs, including TP53 activation, NF-κB signaling, DNA-damage responses, inflammatory signaling, and cellular senescence (14, 46, 51, 60). Although these states may participate in normal epithelial repair, their persistence in fibrotic lungs is associated with failed AT1 differentiation and pathological epithelial remodeling (13, 14, 46, 61), a main observation in our model of AT2-specific mitochondrial damage. Moreover, recent work suggests that KRT8 is not merely a marker of transitional epithelial cells but can contribute functionally to macrophage chemokine expression, macrophage recruitment, and fibrogenesis (13). In this context, the KRT8^+^ and p21^+^ phenotype induced by TFAM loss likely reflects a stressed, aberrant intermediate epithelial state rather than normal AT2-to-AT1 differentiation process. While our data support the emergence of a mitochondrial stress-associated epithelial state, future single-cell and spatial analyses will be required to resolve the spatiotemporal evolution of TFAM-deficient AT2 cells and determine whether they converge with previously described ADI, PATS, DATP, or aberrant basaloid states, or define a distinct mitochondrial dysfunction-associated epithelial state.

Our findings extend to a growing body of work indicating that mitochondrial function is a determinant of alveolar epithelial fate, regenerative capacity, and fibrotic susceptibility, rather than merely a downstream consequence of tissue injury. We and others have shown that mitochondrial activity contributes to acquisition of alveolar epithelial cellular properties, while defective mitophagy, impaired fatty acid oxidation, altered epithelial redox control, and mitochondrial complex I dysfunction can compromise AT2 cell repair, promote aberrant epithelial states, and increase fibrosis susceptibility (18, 30–32, 62, 63). In particular, epithelial deletion of *Ndufs2*, a part of mitochondrial complex I, caused postnatal accumulation of *Krt8*^+^*Cdkn1a*^+^ transitional epithelial cells and impaired alveolar epithelial differentiation through NAD^+^-dependent activation of the integrated stress response, supporting the concept that mitochondrial respiratory dysfunction can directly alter alveolar epithelial cell fate (30). However, the impact of *Ndufs2* in the adult lung, and its potential role on fibrotic lung disease, remains unknown. In this context, TFAM loss represents a distinct and potentially upstream mitochondrial defect because it directly disrupts mitochondrial genome maintenance and broadly suppresses mtDNA-encoded respiratory chain gene expression. Although our Seahorse assays do not resolve the activity of individual electron transport chain complexes, our RNA-seq data suggest that TFAM deficiency is unlikely to represent an isolated complex I defect. Instead, suppression of all 13 mtDNA-encoded protein-coding transcripts predicts broad impairment of mtDNA-dependent respiratory complexes, including complexes I, III, IV, and V, but not complex II. Complex I may be particularly relevant because 7 of the 13 mtDNA-encoded protein-coding genes encode complex I subunits, and complex I is a major regulator of NAD⁺/NADH balance. Thus, the reduced NAD⁺/NADH ratio observed in TFAM-deficient AT2 cells raises the possibility that impaired complex I-linked NAD⁺ regeneration contributes to transitional and senescence-associated remodeling.

TFAM is essential for mtDNA packaging, maintenance, and transcription, and studies in other tissues have shown that TFAM loss can destabilize mtDNA and activate tissue-remodeling pathways (28, 29). In immune cells, T cell-specific *Tfam* deletion induces systemic inflammaging and premature senescence phenotypes (64). In systemic sclerosis fibroblasts, reduced TFAM has been linked to mitochondrial damage and mtDNA release, along with increased susceptibility to fibrotic remodeling (28). In the kidney, *Tfam* deletion has been shown to activate cGAS-STING signaling and contribute to renal fibrosis (29). cGAS-STING is activated by cytosolic DNA and has been extensively studies in innate immune cells to drive inflammatory responses (26, 65). Emerging studies also report impaired cGAS-STING signaling in AT2 cells upon aging (66). Notably, cGAS-STING can further lead to cellular senescence (67), supporting the hypothesis of an aberrant TFAM-cGAS-STING axis driving the transitional AT2 cell phenotype.

Beyond cell-intrinsic epithelial remodeling, TFAM-deficient AT2 cells acquired a broad secretory program capable of reshaping the lung microenvironment. Importantly, this program included multiple secreted mediators previously implicated in pulmonary fibrosis, epithelial stress, senescence-associated signaling, and fibroblast activation, including WISP1/CCN4 (38, 39), and GDF15 (68–70). This finding is consistent with emerging evidence that aberrant epithelial states in fibrosis are central signaling hubs that communicate with fibroblasts, macrophages, and other niche cells (11, 13, 14). Our data extend this concept by identifying epithelial mitochondrial failure as an upstream trigger of profibrotic paracrine signaling. Within this TFAM-regulated secretory program, IGFBP2 emerged as one functionally relevant downstream mediator, as well as potential therapeutic target. IGFBP2 is well-known for its canonical role in regulating IGF bioavailability (71, 72), and has further been described to activate pro-fibrotic pathways, such as FAK/ERK, STAT3, and NF-κB signaling in other tissue-remodeling and disease contexts (73–75). We found increased IGFBP2 transcript and protein expression in multiple independent lung tissue datasets. Functionally, recombinant IGFBP2 promoted fibrotic remodeling, whereas IGFBP2 neutralization attenuated profibrotic signaling induced by TFAM-deficient AT2 cells as well as in human pulmonary fibrosis. Similarly, IGFBP2 was recently shown to promote epithelial barrier disruption and epithelial-mesenchymal transition (EMT)-like remodeling through FAK signaling in an airway epithelial disease model (76). However, our findings differ from a recent study that reported epithelial IGFBP2 to suppress AT2 senescence and protect from bleomycin-induced fibrosis (77). In support of our observations, IGFBP2 has been reported to be elevated in serum and sputum from patients with IPF (40–42). Importantly, our functional data supporting a profibrotic role for IGFBP2 are based on human cells and lung tissue, underscoring potential species-, model-, and context-dependent differences in IGFBP2 biology. Moreover, our data highlight IGFBP2 inhibition as a potential therapeutic target for human lung tissue remodeling in both fibrotic cocktails treated hPCLS as well as IPF tissue *ex vivo*. These data need to be expanded, and downstream analysis is required in future studies to mechanistically elucidate the impact of blocking IGFBP2. Our data provide a strong rationale for IGFB2 being a therapeutic target to address AT2 cell mitochondrial dysfunction.

Previous studies have started to target epithelial mitochondrial function and metabolism as therapeutically relevant axes in fibrotic lung disease. For example, thyroid hormone-mediated restoration of mitochondrial homeostasis in AT2 cells improved mitochondrial bioenergetics and reduced experimental pulmonary fibrosis. Similarly, modulation of key signaling processes, such as PINK1-dependent mitophagy, CYB5R3-mediated redox control, and AT2 fatty acid oxidation has been shown to preserve epithelial mitochondrial fitness and attenuate fibrotic remodeling (16–18, 31, 32). An alternative strategy to target TFAM loss mediated AT2 cell reprogramming is to directly restore TFAM expression or activity. Indeed, a recently described small-molecule TFAM activator has been shown to increase TFAM protein levels, suppress mtDNA stress-associated interferon signaling, and reduce fibrotic marker expression in fibroblasts (78). However, because this compound appears to act on existing functional TFAM protein rather than inducing *TFAM* expression, further studies will be needed to determine its efficacy in TFAM-deficient cells and in relevant *in vivo* models of pulmonary fibrosis. In this context, IGFBP2 neutralization may represent a complementary strategy, analogous to senomorphic approaches that target pathogenic components of the senescence-associated secretory phenotype rather than eliminating senescent cells or broadly disrupting essential cellular functions. By targeting a specific AT2-derived mediator that acts on the surrounding microenvironment, IGFBP2 blockade may attenuate the harmful consequences of TFAM-deficient epithelial secretory remodeling while avoiding direct interference with mitochondrial programs required for healthy cells. Nevertheless, the partial rescue achieved by IGFBP2 blockade indicates that TFAM loss likely engages additional profibrotic mediators and mechanisms, which remain to be defined in future studies.

In conclusion, we identify AT2 cell-intrinsic TFAM deficiency as a driver of transitional AT2 cell emergence and progressive pulmonary fibrosis. Our findings further establish IGFBP2 as a key paracrine mediator linking epithelial mitochondrial dysfunction to profibrotic cell-cell communication and support IGFBP2 blockade as a potential therapeutic strategy for lung fibrosis.

## METHODS

### Sex as a biological variable

Human donor and IPF lung samples were obtained from both sexes. For animal studies, the bleomycin-induced fibrosis model in C57BL/6 mice was performed exclusively in male mice to minimize sex-related variability in injury responses and to maintain consistency with prior bleomycin studies. In contrast, the AT2-specific genetic *Tfam* deletion model included both male and female mice. The study was not designed for sex-stratified comparisons. Therefore, sex was not analyzed as a primary biological variable. Findings are anticipated to be relevant to both sexes.

### Study approval

Human lung specimens, including formalin-fixed paraffin-embedded (FFPE) tissue, and tissue used for the generation of human precision-cut lung slices (PCLS), were obtained from the Human Lung Tissue Biobank of the University of Pittsburgh Division of Pulmonary, Allergy, Critical Care and Sleep Medicine under Institutional Review Board-approved protocols PRO14010265 and CORID #300/#451. Bronchoalveolar lavage fluid (BALF) samples from non-ILD controls and patients with IPF used for cytokine array analyses were obtained from the CPC-M bioArchive at the Comprehensive Pneumology Center (CPC-M, Germany), approved by the local ethics committee of the Ludwig-Maximilians-Universität München, Germany (Ethic vote #19-630), or from the UGMLC Giessen Biobank, Germany (approved by the Ethics Committee of Department 11 at Justus Liebig University Giessen, AZ 58/15). Written informed consent was obtained from all study participants. All animal experiments were approved by the University of Pittsburgh Institutional Animal Care and Use Committee (IACUC 24014329 & 24014453) and performed in accordance with the Guide for the Care and Use of Laboratory Animals.

### Human precision-cut lung slices (hPCLS)

Fresh human lung tissue was processed for hPCLS generation and culture using a previously published workflow (79–81). Briefly, the peripheral lung parenchyma was inflated via a segmental bronchus with 2% low-melting-point agarose in PBS and placed on ice to solidify. Tissue cores were generated using a 10-mm biopsy punch and cut into 300-µm slices with a Compresstome (Precisionary Instruments, VF-310-0Z). Slices were transferred to 24-well plates and cultured in DMEM/F12 supplemented with 0.1% FBS and Penicillin-Streptomycin at 37°C in a humidified incubator with 5% CO_2_. hPCLS were allowed to recover for 12 hours before experimental treatment. Where indicated, hPCLS were exposed to recombinant IGFBP2 with media refreshed daily. Fibrotic responses were assessed by collagen immunostaining as described below.

### Mouse strains and tamoxifen induction

AT2 cell-specific *Tfam* knockout mice were generated by crossing *Sftpc*^CreERT2^ mice (The Jackson Laboratory, stock #028054) with *Tfam*^fl/fl^ mice (The Jackson Laboratory, stock #026123) (22, 82). Two independent tamoxifen-induction cohorts were used in this study. In the diet-induced cohort, experimental groups included wild-type controls lacking Cre recombinase (WT: *Sftpc*^+/+^; *Tfam*^fl/fl^), heterozygous conditional knockout mice (Het: *Sftpc*^CreERT2/+^; *Tfam*^fl/+^) and homozygous conditional knockout mice (KO: *Sftpc*^CreERT2/+^; *Tfam*^fl/fl^). Cre-mediated recombination was induced by tamoxifen-containing diet administered in alternating weekly cycles, with 1 week on tamoxifen diet followed by 1 week on regular diet, for 6 cycles total, corresponding to a 12-week diet intervention. This cohort was used to assess gene-dosage-dependent TFAM reduction, spontaneous lung phenotypes, and responses to saline or bleomycin challenge, as indicated. A separate cohort receiving intraperitoneal tamoxifen cohort used to validate the phenotype with an independent induction strategy. In this cohort, experimental groups included Cre-positive wild-type controls (i.p. WT: *Sftpc*^CreERT2/+^; *Tfam*^+/+^) and AT2 cell-specific Tfam conditional knockout mice (i.p. KO: *Sftpc*^CreERT2/+^; *Tfam*^fl/fl^). Mice received tamoxifen by intraperitoneal injection at 75 mg/kg body weight once daily for 5 consecutive days, followed by two additional booster injections administered on consecutive days every 4 weeks. Littermate controls were used whenever possible.

### Mouse precision-cut lung slices (mPCLS)

*m*PCLS were generated and cultured as described above for hPCLS, with minor modifications (83). Briefly, mice were euthanized, and the pulmonary circulation was perfused with cold PBS. Lungs were inflated through the trachea with 2% low-melting-point agarose in PBS and cooled on ice to solidify. Agarose-inflated lungs were sectioned directly into 300-µm slices using a vibratome (Campden Instruments, 7000smz-2). Slices were transferred to 24-well plates and maintained in DMEM/F12 supplemented with 0.1% FBS and penicillin-streptomycin at 37°C in a humidified incubator with 5% CO2. mPCLS were allowed to recover for 12 hours before experimental treatment.

### Bleomycin-induced pulmonary fibrosis

Where indicated, pulmonary fibrosis was induced by a single intratracheal instillation of bleomycin (2 U/kg). Control mice received intratracheal saline. Mice were harvested and analyzed 14 days after bleomycin administration. Animals were monitored daily, and body weight was recorded throughout the experiment.

### Pulmonary function testing

Respiratory function testing was performed using the flexiVent system (SCIREQ) before terminal sample collection. Mice were anesthetized (with an intraperitoneal ketamine/xylazine mixture, 300 and 100 mg/kg respectively), tracheostomized, and connected to the ventilator (tidal volume of 10 ml/kg, 150 breaths/min, PEEP of 3 cm H_2_O, and FIO_2_ of 21%) initiated. Once on the ventilator, intraperitoneal pancuronium bromide (13 mg/kg) was given for muscle relaxation and to ensure passive ventilation. Readouts of positive end-expiratory pressure and positive inspiratory pressure were recorded in real-time. Baseline data and step-wise pressure-volume curves were obtained after approximately 5 min of equilibration, with no evidence of spontaneous respiratory effort.

### Histology and immunofluorescence imaging

After ligation and resection of the right lung lobes, the left lungs were gravity-fixed by tracheal inflation with 10% neutral-buffered formalin for 10 minutes at room temperature, followed by immersion fixation overnight at 4°C. Lungs were paraffin-embedded and sectioned at 4 μm. Sections were stained with H&E for morphology and Masson’s trichrome for collagen deposition. Fibrosis was quantified as collagen-positive area on Masson’s trichrome-stained sections by investigators blinded to genotype and treatment. For immunofluorescence, paraffin sections were deparaffinized, rehydrated, and subjected to antigen retrieval with Dako Target Retrieval Solution (pH 9). Sections were permeabilized with 0.1% Triton X-100, blocked with 5% normal donkey serum, and incubated with primary antibodies against LAMP3 (Novus Biologicals, Cat# DDX0191P, 1:200), TFAM (Abcam, Cat# ab252432, 1:200), TOM20 (Abcam, Cat# ab186734, 1:200), GFP (Abcam Cat# ab13970, 1:500), IGFBP2 (Invitrogen Cat# PA5-79450, 1:500), KRT8 (DSHB Cat# TROMA-I, 1:500), p21 (Abcam Cat# ab188224, 1:200), Fibronectin (Abcam Cat# ab45688, 1:500), a-SMA (Novus Biologicals, NBP2-33006, 1:300), COL1A1 (Abcam Cat# 19254, 1:500). After washing, sections were incubated with species-appropriate fluorophore-conjugated secondary antibodies and counterstained with DAPI. Slides were mounted in Vector TrueView mounting medium (Vector Cat# H-1700) and imaged on an Olympus IX83 inverted fluorescence microscope using identical acquisition settings across conditions. Image analysis was performed in a blind manner using ImageJ.

### Primary AT2 cell isolation

Primary mouse AT2 (mpAT2) cells were isolated from adult *Tfam*^fl/fl^ mice using an established lung dissociation and epithelial enrichment workflow with minor modifications (83). Briefly, mice were euthanized, and lungs were perfused with cold PBS, followed by enzymatic digestion with dispase (Corning® #354235, 50 U/ml) and mechanical dissociation. Single-cell suspensions were filtered through 100-, 70-, and 40-µm strainers, subjected to red blood cell lysis as needed. Negative selection of fibroblasts was performed by adherence to non-coated plastic plates. Macrophages and white blood cells were depleted using CD45- and endothelial cells were depleted with CD31-specific magnetic beads (Miltenyi Biotec, Bergisch Gladbach, Germany) according to the manufacturer’s instructions.

### In vitro Cre recombinase-mediated Tfam deletion

Purified mpAT2 cells were plated on well plates and maintained in DMEM/F12-based medium supplemented with 10% FBS, Penicillin-Streptomycin. To induce *Tfam* deletion, cells were infected with adenoviral Cre recombinase (AdCre+; MOI 20; U of Iowa-5 Ad5CMVCre) for 5 h in DMEM/F12-based medium, while control cultures received a matched control adenovirus (AdCre−; MOI 20; U of Iowa-272 #Ad5CMVempty) under identical conditions. After infection, virus-containing media were replaced with fresh culture medium. Cells were re-infected on day 3, and downstream analyses were performed on day 7. Validation of *Tfam* deletion. Deletion efficiency was confirmed by qPCR for *Tfam* transcript levels and TFAM immunofluorescence in cultured mpAT2 cells. Where indicated, mitochondrial perturbation was assessed by mtDNA copy number relative to genome DNA copy number.

### Conditioned medium (CM) collection from primary AT2 cultures

Conditioned media were generated from primary mpAT2 cultures following adenoviral Cre-mediated *Tfam* deletion. Briefly, mpAT2 cells were infected with AdCre− or AdCre+ as described above. On day 6 after the initial infection, cultures were washed once with prewarmed PBS and the medium was replaced with phenol red-free DMEM/F12-based medium. After 24 h, conditioned media were collected and clarified by centrifugation at 14,000g for 15 min at 4°C to remove cellular debris. The supernatant was transferred to a fresh tube and used immediately for downstream assays or aliquoted and stored at −80°C until use.

### Fibroblast migration (wound-scratch) assay

CCL-206 fibroblasts were seeded at 3 × 10^4^ cells per well into removable 2-well culture inserts (ibidi, Cat# 80242) placed in 24-well plates to generate a defined cell-free gap. After cells reached confluence, cultures were serum-starved for 24 h in medium containing 0.1% FBS. Inserts were then removed to initiate migration, and cells were immediately treated with conditioned media collected from AdCre− or AdCre+ mpAT2 cultures. Plates were placed in a Cytation live-cell imaging system (Agilent/BioTek), and phase-contrast images were acquired hourly for 24 h. Migration was quantified as percent wound closure by measuring the remaining gap area over time relative to time 0 using Gen5.

### Cytokine array profiling

Cytokine profiles in mouse AT2 conditioned media and human bronchoalveolar lavage (BAL) fluid were assessed using Proteome Profiler XL cytokine arrays (R&D Systems; Mouse XL ARY028 and Human XL ARY022B) according to the manufacturer’s instructions. Clarified conditioned media or BALF were normalized by volume and incubated with the array membranes overnight at 4°C. Membranes were washed, incubated with the supplied detection antibody cocktail followed by streptavidin-HRP, and developed using a chemiluminescent substrate. Signals were captured on a ChemiDoc imaging system using identical exposure settings across conditions. BAL samples were clarified by centrifugation to remove cells and debris and were applied to membranes neat or diluted as recommended by the manufacturer. Spot intensities (duplicate spots per analyte) were quantified in ImageJ after local background subtraction and normalized to internal reference/positive control spots on each membrane.

### ELISA assay

Previously collected and archived conditioned medium from control cocktail (CC) and fibrotic cocktail (FC) treated human PCLS was analyzed for IGFBP2 abundance using a human IGFBP-2 DuoSet ELISA kit (R&D Systems, #DY674) according to the manufacturer’s instructions. Samples and standards were assayed in duplicate. Absorbance was measured at 450 nm with wavelength correction at 540 nm. Paired CC and FC samples from the same donor lung were analyzed in parallel.

### Bulk RNA sequencing and bioinformatic analysis

Bulk RNA sequencing was performed on snap-frozen whole mouse lung tissue and on primary mpAT2 cultures following adenoviral Cre-mediated *Tfam* deletion (AdCre− vs AdCre+). Total RNA was isolated using the RNeasy Plus Mini Kit (Qiagen, Cat# 74104), and RNA integrity was assessed by Agilent Bioanalyzer (RNA Nano 6000). A total of 1 μg RNA per sample was submitted to Novogene for library preparation and sequencing. Briefly, mRNA was purified from total RNA by poly(A) selection using poly-T oligo-attached magnetic beads, fragmented, and reverse-transcribed to generate first-strand cDNA using random hexamer primers, followed by second-strand cDNA synthesis. Libraries were generated using the NEBNext Ultra RNA Library Prep Kit for Illumina (non-strand-specific), including end repair, A-tailing, adaptor ligation, size selection, PCR amplification, and purification. Pooled libraries were sequenced on an Illumina NovaSeq 6000 platform (S4 flow cell) to generate paired-end 150-bp reads (PE150), targeting approximately 20 million read pairs (∼6 Gb) per sample. Primary data processing followed a standard pipeline. FASTQ files were quality-filtered using fastp to remove adaptor sequences, poly-N reads, and low-quality reads, and to compute read-quality metrics (including Q20/Q30 and GC content). Clean reads were aligned to the mouse reference genome (GRCm39) using HISAT2 (v2.2.1). Gene-level read counts were generated using featureCounts (Subread v2.0.1). Differential gene expression was performed using DESeq2 with Benjamini-Hochberg correction; adjusted P < 0.05 was considered significant.

### Transmission electron microscopy (TEM)

Cells were fixed in cold 2.5% glutaraldehyde in PBS (pH 7.3) and pelleted by centrifugation at 300g. Cell pellets were rinsed in PBS, post-fixed in 1% osmium tetroxide containing 1% potassium ferricyanide, dehydrated through a graded ethanol series, and embedded in Poly/Bed 812 resin (Glauert formulation). Section of 300 nm were cut using a Leica Reichart Ultracut ultramicrotome, stained with 0.5% toluidine blue in 1% sodium borate, and examined by light microscopy for sample orientation. Ultrathin sections (65 nm) were then stained with Uranyless and Reynolds’ lead citrate and imaged using a JEOL 1400 Plus transmission electron microscope equipped with a side-mount Nikon 50M digital camera.

### Seahorse extracellular flux analysis

Mitochondrial function was assessed in *Tfam*-deleted and control AT2 cells using the Seahorse XF Cell Mito Stress Test on an XF96 Extracellular Flux Analyzer (Agilent). Cells were detached, seeded into poly-L-lysine-coated Seahorse XF96 assay plates, and incubated for 4 hours to allow attachment. Cells were then washed and incubated in Seahorse XF assay medium (pH 7.4) before oxygen consumption rate (OCR) measurements according to the manufacturer’s protocol. Following three baseline measurements, oligomycin (2.5 μM), FCCP (1 μM), and rotenone/antimycin A (0.5 μM each) were sequentially injected. Real-time OCR measurements were used to determine basal respiration, maximal respiration, ATP-linked respiration, and proton leak. OCR values were normalized to total cellular mass quantified using the CyQUANT assay (Molecular Probes, Thermo Fisher Scientific) according to the manufacturer’s instructions.

### Statistics

For pairwise analyses, significance was assessed using an unpaired, 2-tailed parametric t-test. For experiments involving three or more groups, differences were evaluated by 2-way ANOVA followed by Tukey’s adjustment for multiple comparisons. Statistical significance was defined as P < 0.05. Results are reported as mean ± SEM.

### Data availability

Bulk RNA-seq data generated in this study have been deposited in the NCBI Gene Expression Omnibus (GEO) under accession numbers GSE315898 (whole-lung RNA-seq from AT2-specific *Tfam* knockout mice) and GSE315897 (*in vitro* Cre recombinase-mediated *Tfam* deletion). Codes for data processing and analysis were reported in the GitHub repository (https://github.com/QianjiangHu/TFAM_Fibrosis_2026).

## ACKNOWLEDGMENTS

We thank the members of the Königshoff laboratory for helpful discussions and technical support. We also thank members of the Bueno and Kauffman laboratories for their technical assistance with the generation and preparation of rodent cohorts. We gratefully acknowledge the Human Lung Tissue Biobank of the University of Pittsburgh Division of Pulmonary, Allergy, Critical Care, and Sleep Medicine for providing human lung specimens, including formalin-fixed paraffin-embedded (FFPE) tissue, and tissue used for generation of human precision-cut lung slices. We also gratefully acknowledge the provision of bronchoalveolar lavage fluid (BALF) samples and associated clinical data from the CPC-M bioArchive at the Comprehensive Pneumology Center Munich and its partners at the LMU University Hospital, Ludwig-Maximilians-Universität München and the Asklepios Biobank Gauting, and from the UGMLC Giessen Biobank. We thank the patients and their families for their support.

This work was supported by the NIH Common Fund U54AG075931, the Three Lakes Foundation, and NIH P50AR080612 to MK, and by NIH R01HL146519 to OE. ML was supported by the Deutsche Forschungsgemeinschaft (DFG, German Research Foundation; 512453064 and 54210281), the von Behring Röntgen Foundation (71_0011), the Hessisches Ministerium für Wissenschaft und Forschung, Kunst und Kultur (LOEWE Habitat), and the German Center for Lung Research (DZL). CRK was supported by the Burroughs Wellcome Fund Career Award for Medical Scientists. This work also used resources from the University of Pittsburgh Center for Research Computing and Data, RRID: SCR_022735, including the HTC cluster supported by NIH award S10OD028483. Transmission electron microscopy was performed at the Center for Biologic Imaging, University of Pittsburgh, using instrumentation supported by NIH grant S10OD016236.

