## supplemental figures for "Alveolar Epithelial Cell Loss of the Mitochondrial Regulator TFAM Drives Progressive Lung Fibrosis"

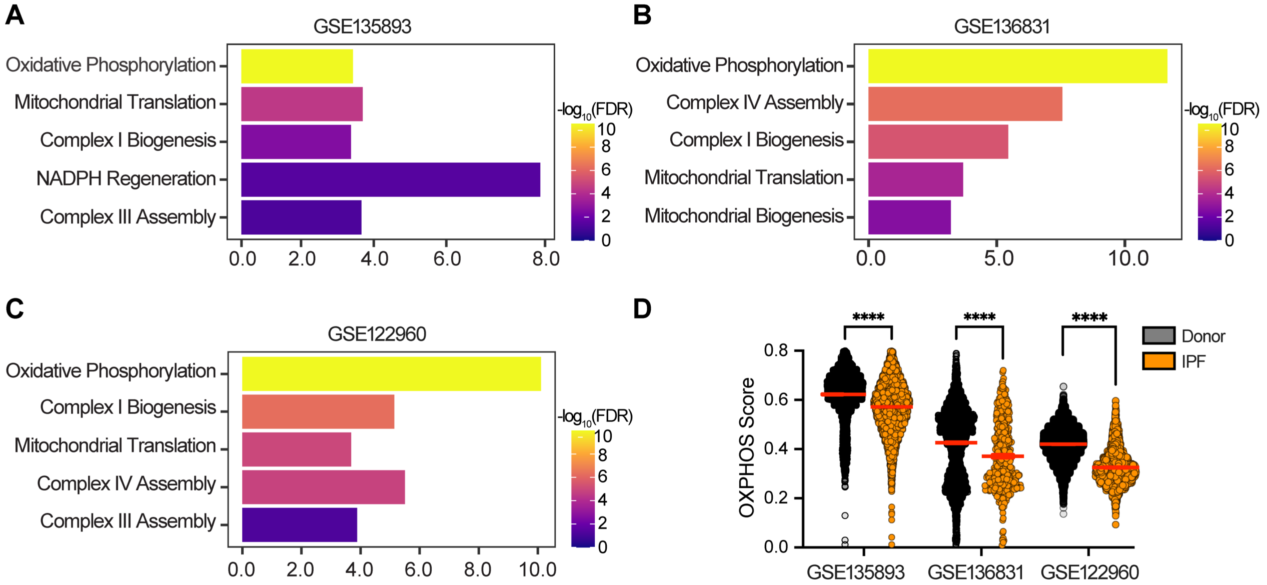


**Supplemental Figure 1. AT2 cells from IPF lungs exhibit mitochondrial dysfunction.**

(A-C) Gene set enrichment analysis of AT2 cells from IPF lungs versus donor controls in three independent datasets (GSE135893, GSE136831, and GSE122960) shows significant enrichment of mitochondrial dysfunction related pathways in IPF. (D) Analysis of oxidative phosphorylation (OXPHOS) activity scores across the same datasets shows reduced OXPHOS signaling in AT2 cells from IPF lungs compared with donor controls; **** $P < 0.0001$.

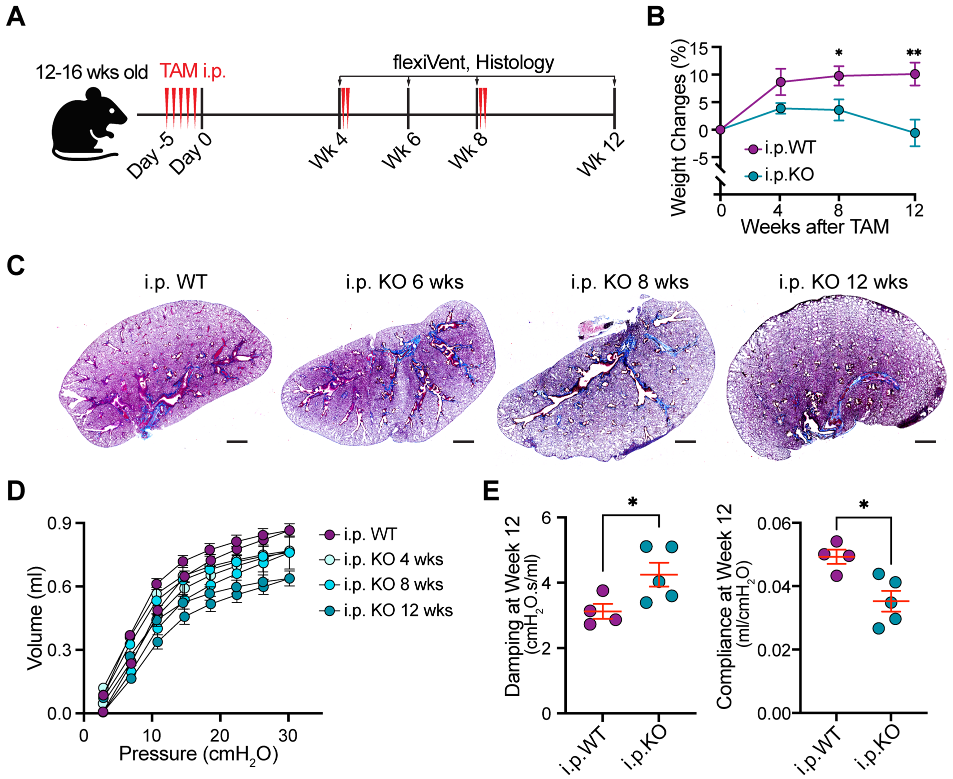


**Supplemental Figure 2. Intraperitoneal tamoxifen administration in AT2 cell-specific *Tfam* conditional knockout mice (i.p. KO) induces progressive lung fibrosis.**

(A) Experimental schematic showing tamoxifen (TAM) delivery by intraperitoneal (i.p.) injection (75 mg/kg) in wild-type controls (i.p. WT: *Sftpc*$^{CreERT2/+}$; *Tfam*$^{+/+}$) and AT2-specific *Tfam* conditional knockout mice (i.p. KO: *Sftpc*$^{CreERT2/+}$; *Tfam*$^{fl/fl}$). (B) Longitudinal body-weight trajectories shows progressive weight loss in i.p. KO mice relative to i.p. WT controls after tamoxifen administration. n = 6 per group. (C) Representative Masson's trichrome-stained lung sections from i.p. WT and i.p. KO mice at 6, 8, and 12 weeks post-tamoxifen injection show progressive fibrotic remodeling and collagen deposition in i.p. KO lungs. Scale bars, 1 mm. (D) Pressure-volume loop analysis at 12 weeks shows impaired lung mechanics in i.p. KO mice compared with i.p. WT controls. n = 5 per group. (E) Lung function measurements 12 weeks after tamoxifen treatment show increased tissue damping and reduced static compliance in i.p. KO mice compared with i.p. WT controls. n = 4 for WT and n = 5 for KO group. Data are presented as mean ± SEM; * $P < 0.05$, ** $P < 0.01$.

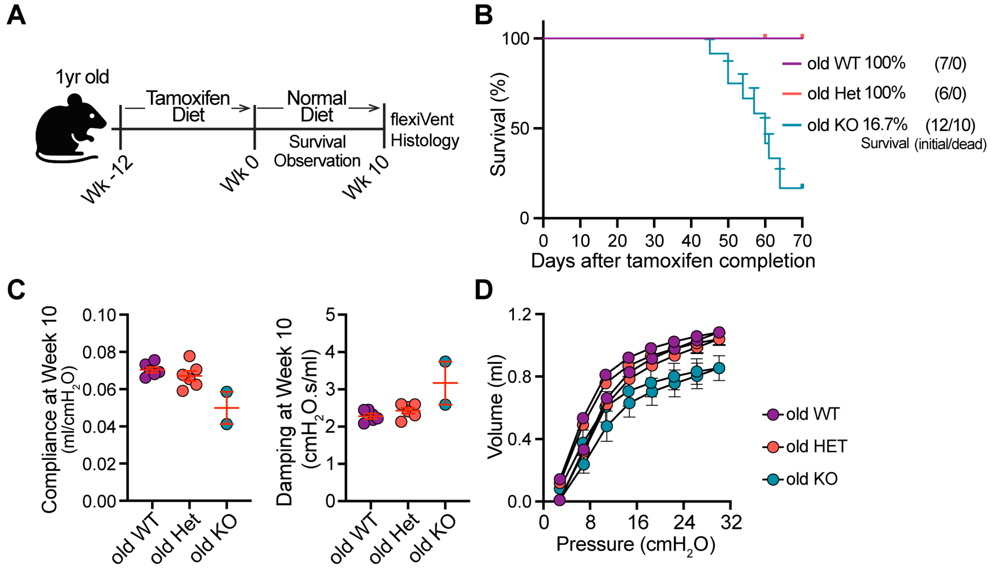


**Supplemental Figure 3. AT2-specific *Tfam* deletion initiated in 12-month-old mice increases mortality and impairs lung mechanics.**

(A) Schematic of the experimental design for tamoxifen-induced, AT2 cell-specific *Tfam* deletion initiated in 12-month-old mice. Mice were maintained on tamoxifen diet for 12 weeks, followed by normal diet for 10 weeks. Week 0 indicates the end of tamoxifen administration and the start of survival monitoring. (B) Kaplan-Meier survival curves show reduced survival of old KO mice (16.7%, n = 12) compared with old Het (100%, n = 6) and old WT (100%, n = 7) controls. Day 0 in panel B corresponds to Week 0 in panel A. (C) Lung mechanics measurements show reduced static compliance and increased tissue damping in the surviving old KO mice compared with old Het and old controls. (D) Pressure-volume loop analysis further confirms impaired lung mechanics in surviving old KO mice compared with old Het and old WT controls. n values are indicated in the graphs.

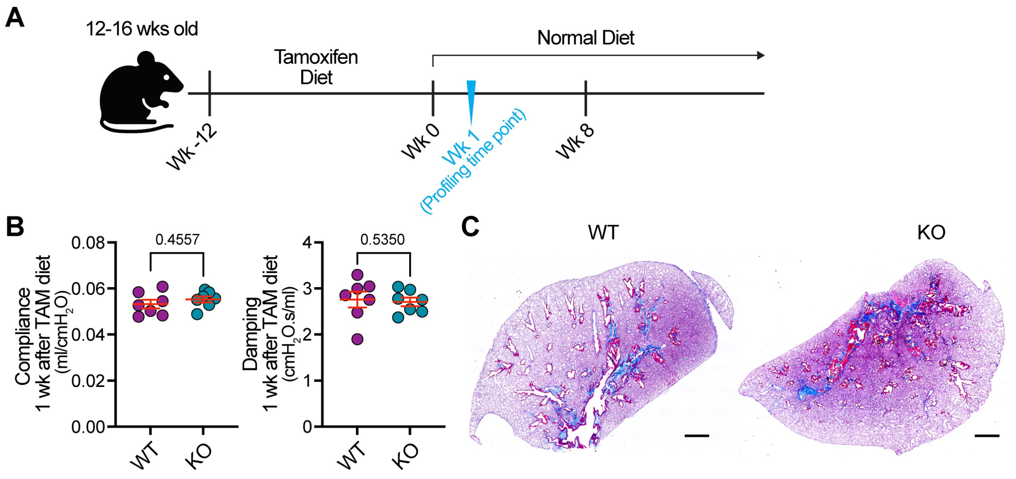


**Supplemental Figure 4. Pre-fibrotic characterization of AT2-specific *Tfam*-KO mice.**

(A) Experimental schematic. AT2-specific *Tfam* deletion was induced in adult mice by tamoxifen diet, and lungs were analyzed 1 week after completion of the 12-week tamoxifen diet, corresponding to a pre-fibrotic profiling time point. (B) Lung function measurements (static compliance and tissue damping) in WT and AT2-specific *Tfam*-KO mice at the pro-fibrotic profiling time point show preserved global mechanics, indicating absence of advanced fibrosis. n = 7 per group. (C) Representative Masson's trichrome-stained lung sections from WT and AT2-*Tfam* KO mice confirm minimal collagen deposition at this stage. n = 7 per group. Scale bars, 1 mm.

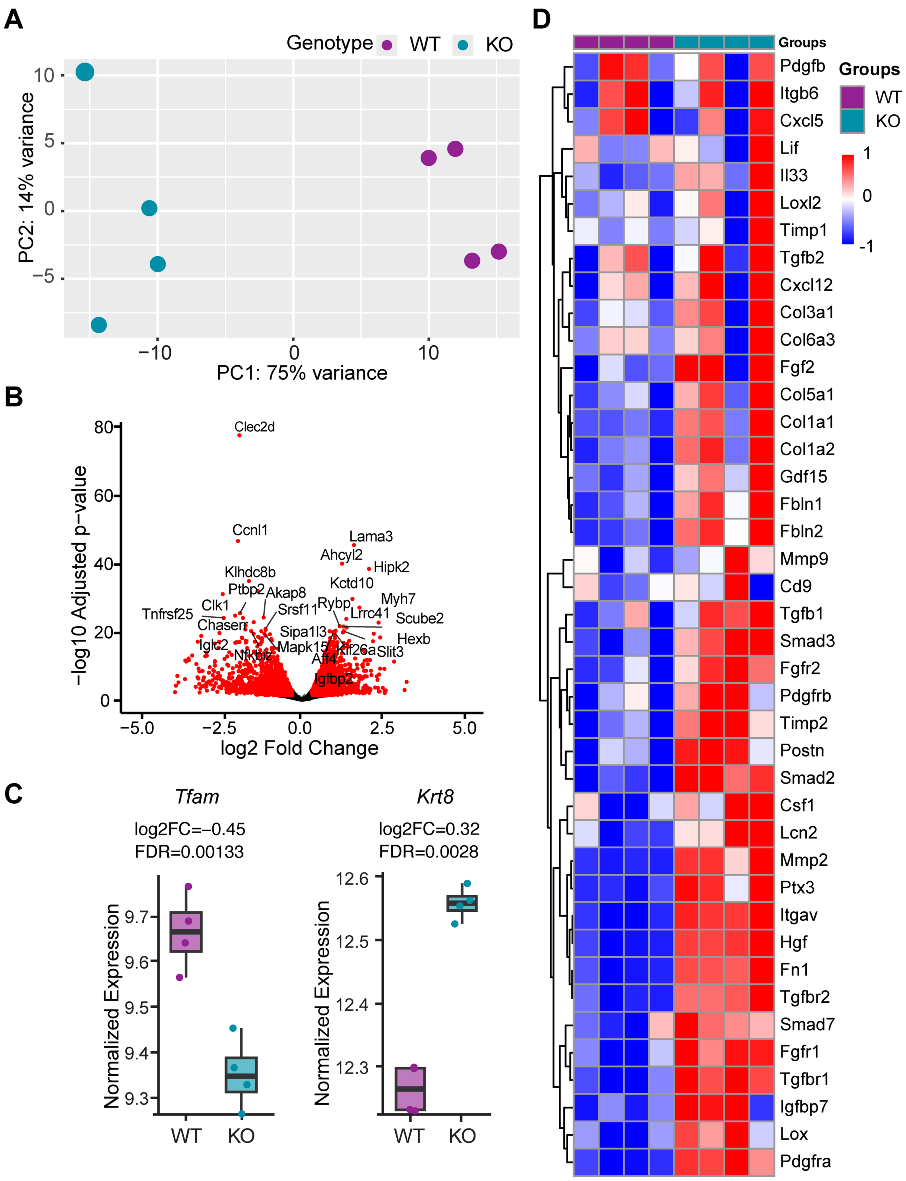


**Supplemental Figure 5. Bulk RNA-seq of whole lungs from WT and AT2-*Tfam* KO mice at pre-fibrotic stage.**

(A) Principal component analysis (PCA) of whole-lung RNA-seq shows clear separation of wild-type (WT) and AT2-*Tfam* knockout (AT2-*Tfam* KO) samples (n = 4 per group), indicating a robust genotype-dependent transcriptional signature. (B) Volcano plot depicting differential gene expression between AT2-*Tfam* KO and WT lungs; significantly altered transcripts (adjusted $P < 0.05$) are highlighted in red, with selected genes labeled. (C) Box plots of normalized expression values for *Tfam* and *Krt8* gene expression in WT and AT2-*Tfam* KO lungs. (D) Heat map of selected differentially expressed genes (DEGs) illustrates consistent activation of pro-fibrotic signaling in AT2-*Tfam* KO lungs.

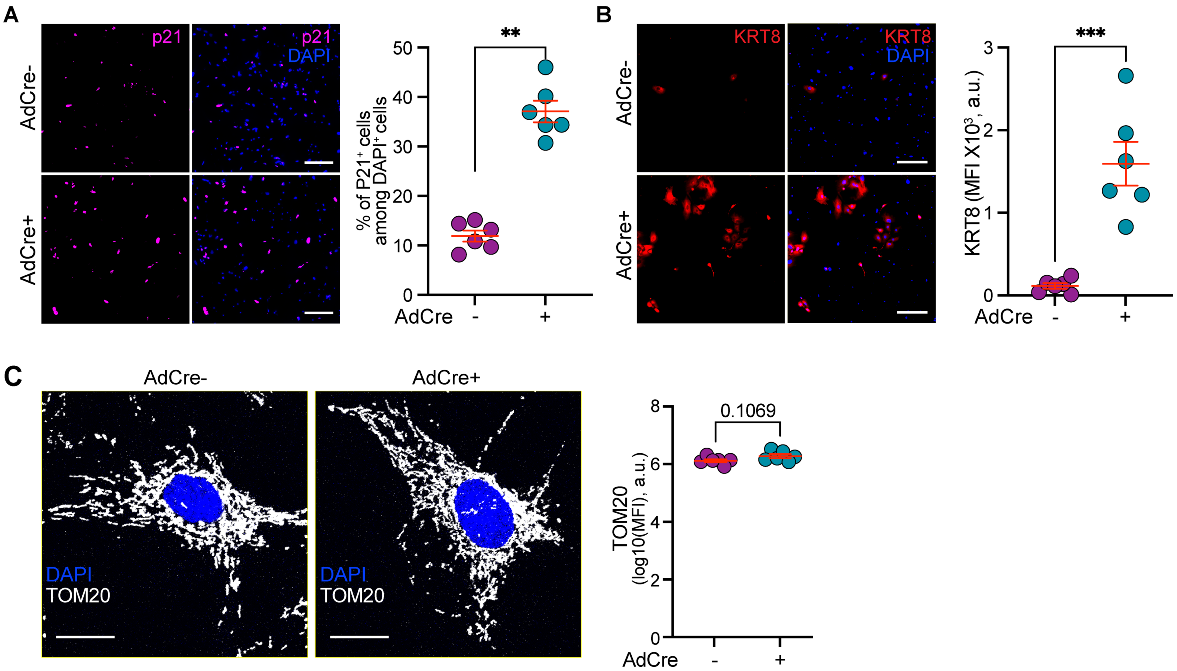


**Supplemental Figure 6. Supporting analysis of TFAM-deficient AT2 cells *in vitro*.**

(A) Representative immunofluorescence images and quantification showing increased p21$^{+}$ AT2 cells following *Tfam* deletion (p21, magenta; DAPI, blue; quantified as percentage of p21$^{+}$ cells relative to total DAPI nuclei). Scale bars, 200 μm. (B) Representative immunofluorescence images and quantification demonstrating accumulation of KRT8$^{+}$ transitional cells in AdCre+ cultures (KRT8, red; DAPI, blue). Scale bars, 200 μm. (C) Immunofluorescence staining showed comparable overall TOM20 abundance in control (AdCre−) and *Tfam*-deleted (AdCre+) AT2 cells, with quantification of TOM20 mean fluorescence intensity (MFI; a.u.) shown at right. Scale bars, 20 μm. Unless otherwise indicated, n = 6 per group. Data are presented as mean ± SEM; ** $P < 0.01$, **** $P < 0.0001$.

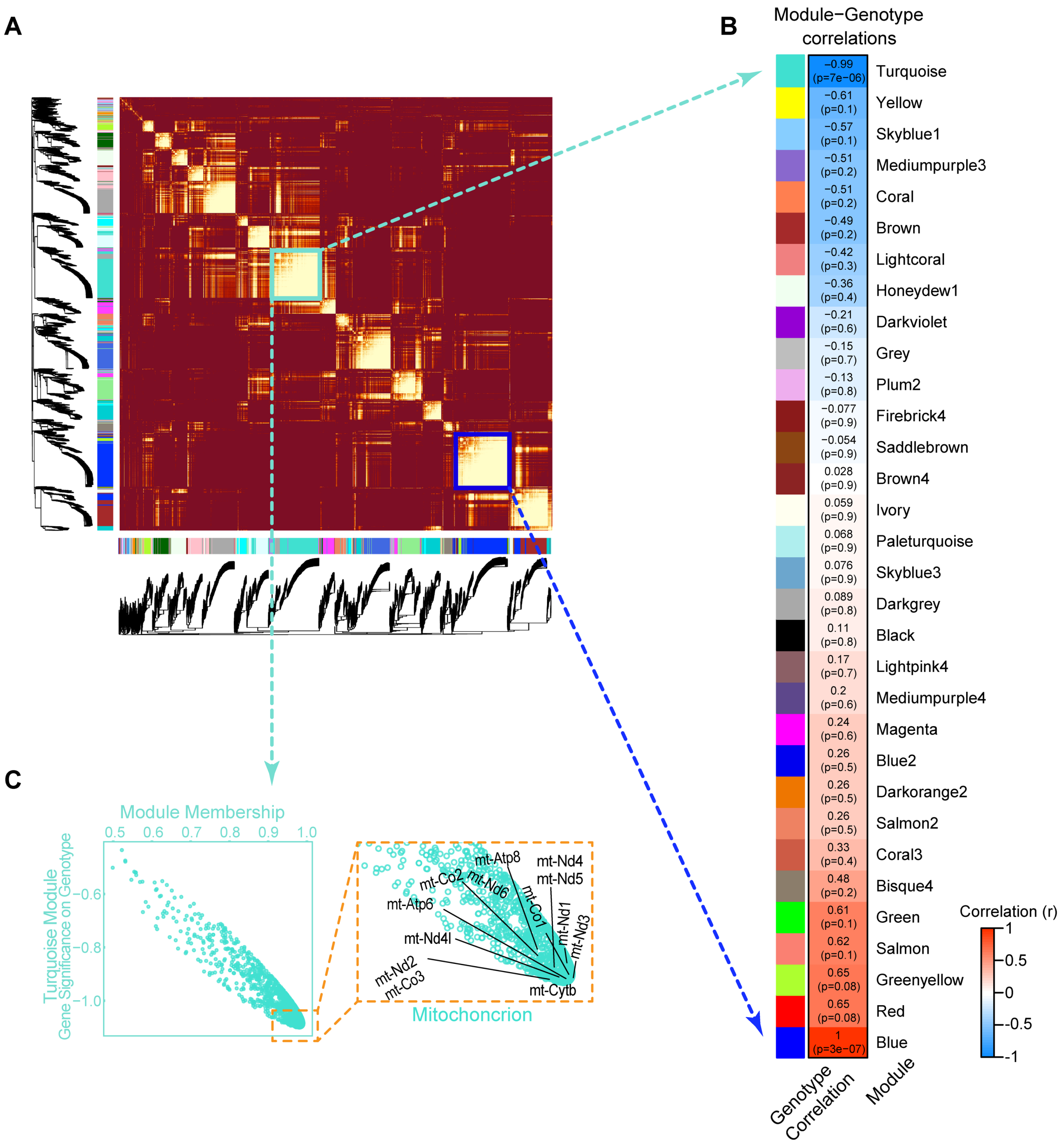


**Supplemental Figure 7. WGCNA identifies co-expression modules associated with *Tfam* deletion in primary AT2 cells.**

(A) Weighted gene co-expression network analysis (WGCNA) was performed using bulk RNA-seq data from control (AdCre−) and *Tfam*-deleted (AdCre+) primary mouse AT2 cells. The heatmap shows the topological overlap matrix (TOM) of expressed genes ordered by average-linkage hierarchical clustering. Warmer colors indicate higher topological overlap and stronger gene co-expression. Dendrograms and adjacent color bars indicate modules identified by dynamic tree cutting. Turquoise- and blue-outlined regions highlight modules with strong internal connectivity. (B) Module-genotype correlations showing the relationship between each module eigengene and *Tfam* deletion status. Colors indicate the direction and strength of the Pearson correlation coefficient, with corresponding p values shown for each module. (C) Scatter plot showing gene significance for *Tfam* deletion versus module membership within the turquoise module, which was strongly negatively correlated with *Tfam* genotype. The inset highlights mtDNA-encoded protein-coding genes.

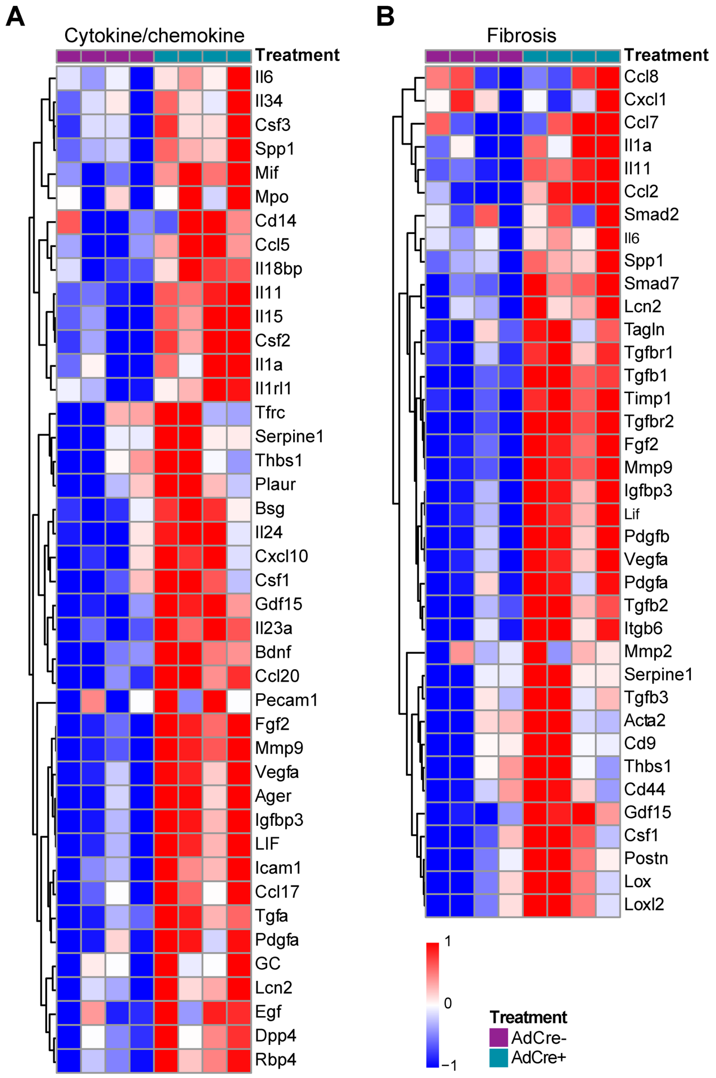


**Supplemental Figure 8. *Tfam* deletion in AT2 cells induces a unique profibrotic transcriptional programs *in vitro*.**

Heatmaps of bulk RNA-seq data showing the expression of selected genes in control (AdCre−) and *Tfam*-deleted (AdCre+) AT2 cells. (A) Selected DEGs annotated to cytokine/chemokine activity (GO:0005125). (B) Selected DEGs annotated to the WikiPathways Lung Fibrosis pathway (WP3624) and/or Reactome Extracellular Matrix Organization pathway (R-HSA-1474244). Each column represents one biological replicate. n = 4 per group.

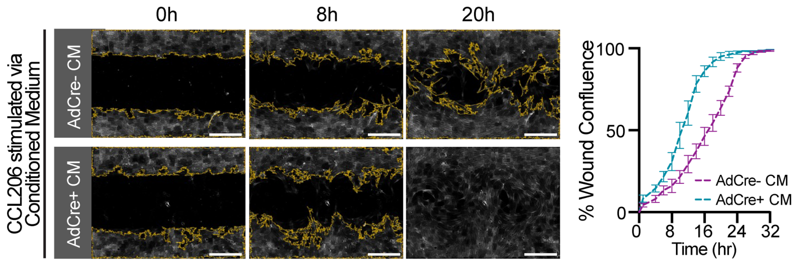


**Supplemental Figure 9. TFAM-deficient AT2 cell secretomes promotes fibroblast migration and collagen expression.**

Conditioned medium from AdCre+ AT2 cells accelerates lung fibroblast (CCL206) migration in wound-scratch assays compared with conditioned medium from AdCre− cells, as shown by representative images and quantified by wound confluence over time. n = 4 per group. Scale bar, 300 μm.

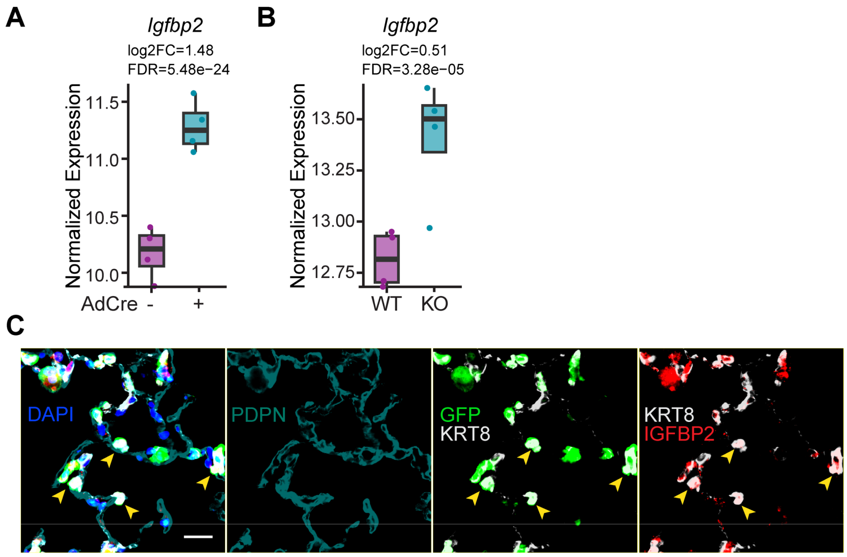


**Supplemental Figure 10. IGFBP2 is upregulated in TFAM-deficient AT2 cells and Krt8⁺ transitional AT2-lineage cells.**

(A) Bulk RNA-seq of primary AT2 cells isolated from *Tfam*$^{fl/fl}$ mice and transduced *in vitro* with adenoviral Cre recombinase (AdCre+) shows increased *Igfbp2* expression compared with control (AdCre−) AT2 cultures. n = 4 per group. (B) Bulk RNA-seq of whole lungs from WT and AT2-specific *Tfam* knockout mice shows increased *Igfbp2* expression in *Tfam* KO lungs. n = 4 per group. (C) In AT2 lineage-tracing *Tfam* knockout mice (Sftpc-CreERT2; mTmG; *Tfam*$^{fl/fl}$), GFP-labeled AT2-lineage cells (green) co-express KRT8 (white) and IGFBP2 (red). Yellow arrowheads indicate GFP$^{+}$KRT8$^{+}$IGFBP2$^{+}$ cells adjacent to PDPN$^{+}$ AT1 cells (cyan). n = 6 per group. Scale bar, 20 μm.
